# High-throughput tagging of endogenous loci for rapid characterization of protein function

**DOI:** 10.1101/2022.11.16.516691

**Authors:** Joonwon Kim, Alexander F. Kratz, Shiye Chen, Jenny Sheng, Liudeng Zhang, Brijesh Kumar Singh, Alejandro Chavez

**Author notes:** Current address: Department of Biomedical Sciences, Cedars-Sinai Medical Center, Los Angeles, CA, 90048, USA.

## Abstract

To facilitate the interrogation of protein function at scale, we have developed High-throughput Insertion of Tags Across the Genome (HITAG). HITAG enables users to rapidly produce libraries of cells, each with a different protein of interest C-terminally tagged is based on a modified strategy for performing Cas9-based targeted insertions, coupled with an improved approach for selecting properly tagged lines. Analysis of the resulting clones generated by HITAG reveals high tagging specificity with the majority of tagging events being indel free. Using HITAG, we fuse mCherry to a set of 167 stress granule-associated proteins and elucidate the features which drive a subset of proteins to strongly accumulate within these transient RNA-protein granules.

## Introduction

To obtain an accurate working model of the cell, we must first understand the dynamic behavior and interaction partners for each of the thousands of proteins within it. A powerful approach to interrogate protein function is through the use of protein tags which are often appended to the C-terminus of a protein of interest, to minimize their influence on protein folding and localization^1,2^. When fused to a protein, these tags enable a myriad of studies such as the *in vivo* examination of protein localization (e.g. fluorescent protein tag), the affinity purification of a protein and its interaction partners from cells and tissues (e.g. FLAG epitope tag), or the rapid destruction of a target protein (e.g. FKBP-DD small molecule regulated degron tag)^3–6^.

As CRISPR/Cas9 has simplified the modification of mammalian genomes, there has been growing interest in tagging all human proteins at their endogenous loci to facilitate the comprehensive mapping of protein behavior^7^. To achieve this goal, groups have relied on homologous recombination to insert tags at the C-terminus of target genes or used non-homologous end joining (NHEJ) to insert synthetic exons containing protein tags into the introns of numerous target genes^7,8^. While powerful, these approaches still involve significant amounts of labor for each line generated or can perturb protein function due to the tag being inserted into the middle of the protein, respectively.

To fulfill the need for a scalable method of gene tagging that would minimally perturb protein function, we developed <u>H</u>igh-throughput Insertion of <u>T</u>ags <u>A</u>cross the <u>G</u>enome (HITAG). HITAG uses Cas9 in combination with non-homologous end joining to insert protein tags into the C-terminus of target genes. Key to the HITAG approach is that the process occurs within a mixed pool of cells wherein, at the end of the procedure, each cell harbors a distinct protein that is C-terminally tagged. In analyzing the insertion events mediated by HITAG, over 70% were found to be “perfect” fusions between the tag and the target gene without the insertion or deletion of additional bases. To enable HITAG, we developed a modified approach to NHEJ-based insertions and an optimized selection marker which dramatically improved the quality of tagged libraries obtained. Using HITAG, a set of 167 stress granule (SG)-associated genes were fused at their endogenous locus with mCherry and the features driving some of these proteins to strongly accumulate within SGs were explored. These results demonstrate the utility of HITAG and provide a means for the scaled interrogation of protein function and behavior across cell types and cellular states.

## Results

### Generating pools of tagged cells using Cas9+NHEJ

To develop HITAG, we began with a previously reported method of NHEJ-based endogenous gene tagging termed CRISPaint^9^. In CRISPaint, a donor plasmid containing the tag to be inserted into the genome is transfected into cells. Along with the donor plasmid, 3 other plasmids containing Cas9, a gRNA against the C-terminus of the gene to be tagged (target-gRNA), and a gRNA against the donor plasmid (donor-gRNA) are also delivered (**Supplementary** Figure 1). Once all plasmids are inside the cell, Cas9 complexes with the target-gRNA and the donor-gRNA to cut the target gene and the donor plasmid, respectively. The cleaved donor plasmid can then become inserted into the endogenous locus via NHEJ. If the tag gets knocked in-frame to the gene of interest, it will also lead to the expression of a downstream drug-resistant marker, enabling the facile enrichment of properly tagged cells by applying drug selection.

While powerful, CRISPaint is limited in that it can only be applied to one target gene at a time, making it laborious to study a large number of genes. To adapt this system into one suitable for rapidly generating libraries of tagged lines, the requirement for performing independent transfections for each target gene needed to be removed. To address this bottleneck, a mixture of target-gRNAs was packaged into lentiviruses and delivered to cells at a low multiplicity of infection (MOI ∼0.1) to ensure that on average, each cell integrates at most a single target-gRNA into its genome. This pool of cells was then transfected *en masse* with the remaining components required for tagging (Cas9, donor plasmid, donor-gRNA), thereby enabling each cell to uniquely tag the gene to which its integrated target-gRNA is directed (**Figure 1a**). Finally, to enrich properly tagged cells, a round of drug selection was applied.

**Figure 1.**
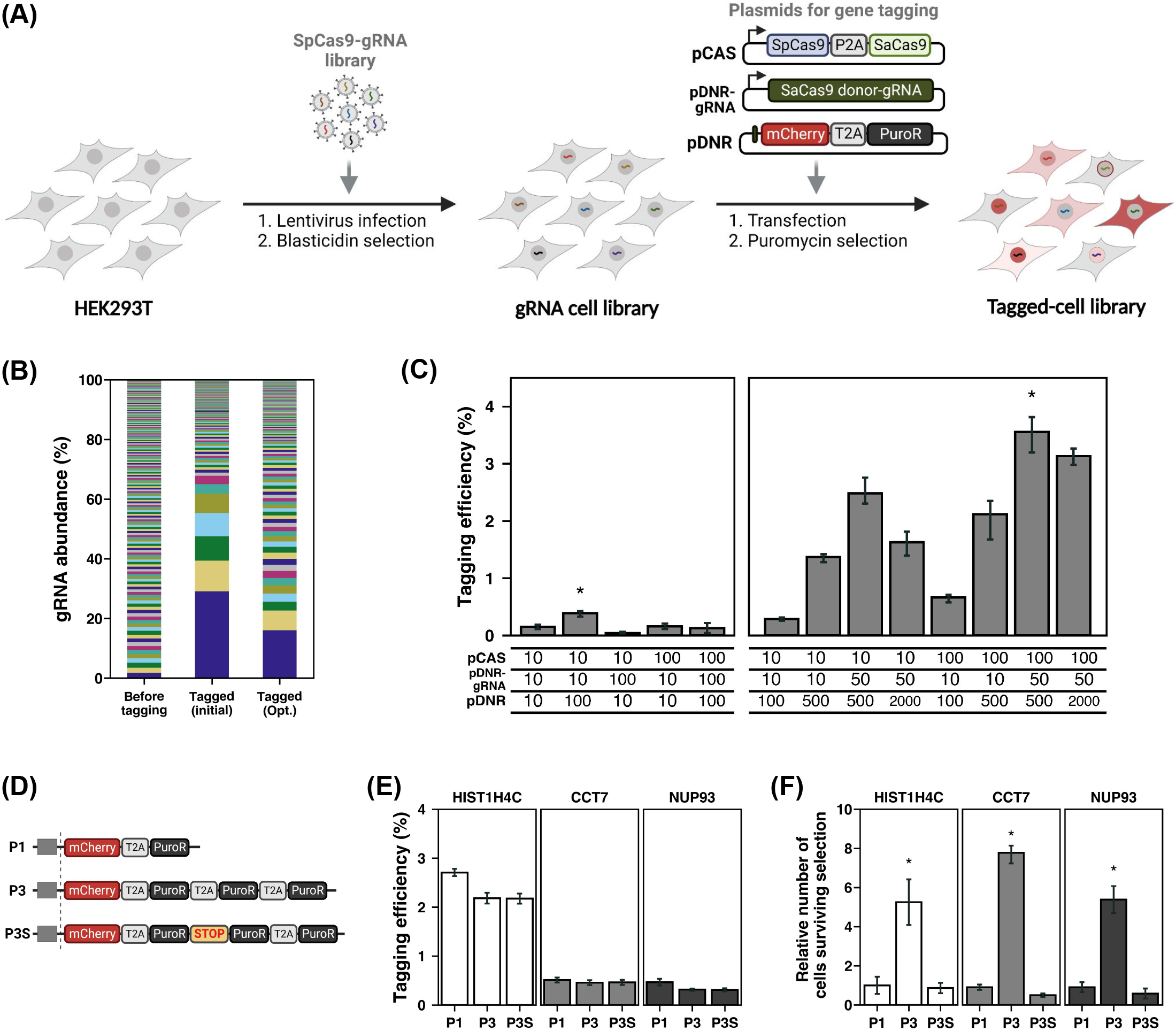
Development of HITAG and properties of the generated lines (a) Schematic summary of HITAG. (b) Distribution of gRNAs within the population of cells before tagging and after tagging and drug selection with the initial versus optimized HITAG approach. Each colored bar represents the abundance of one gRNA within the population. (c) Tagging efficiency before drug selection as a function of different ng amounts of pCAS, pDNR-gRNA, and pDNR plasmid. Data shown are from three biological replicates (independent transfections), the error bars indicate ± the standard deviation. The optimal condition with an asterisk over it showed a significant difference (two sample t-test, * = p < 0.05) to all other conditions tested. (d) Different donor plasmid constructs with one or three copies of the selection marker. P1: a single copy of puroR, P3: three copies of puroR, P3S: three copies of puroR but a stop codon is placed after the first puroR copy. (e) Comparing tagging efficiency when using the donor plasmids with varying numbers of the puromycin resistance gene. (f) The relative number of cells surviving puromycin selection when different donor plasmids with one or three copies of the selection marker were used, data are normalized to the P1 condition. Data shown are from three biological replicates (independent transfections), the error bars indicate ± the standard deviation, the donor with an asterisk over it showed a significant difference (two sample t-test, * = p < 0.05) to all other donors tested across targets.

In optimizing this high-throughput approach, we sought to avoid competition between the target-gRNA and donor-gRNA for complexing with Cas9, since if competition did occur, it would interfere with either cleavage of the endogenous locus or the donor plasmid and in doing so decrease the rate of knock-in^10,11^. To achieve this goal, two orthogonal Cas9 proteins, SpCas9 and SaCas9, were employed, each of which interacts with a unique gRNA scaffold. In our design, SpCas9 is used to cleave the endogenous target gene and SaCas9 is directed to linearize the donor plasmid (**Supplementary** Figure 2). Another benefit of using two orthogonal Cas9 proteins is that properly tagged genes will not be recut, which can occur when only using a single Cas9 protein should the target gene and the donor plasmid have a similar sequence proximal to their PAM sites (**Supplementary** Figure 3). Finally, to reduce off-target insertions which can occur if the donor-gRNA were to inappropriately cleave the genome, several different SaCas9 donor-gRNAs were examined.

From this analysis, a donor-gRNA that showed high tagging efficiency and specificity was identified and used throughout the remainder of the study (**Supplementary** Figure 4). As a “perfect” fusion, without the loss or addition of bases, is the most common repair outcome from tagging, we developed three donor plasmids that upon fusion to the target gene will place the tag in the appropriate translational reading frame (**Supplementary** Figure 5)^9^. This design enables users to achieve precise tagging by selecting an optimal donor plasmid depending upon the location within the C-terminal codon where the target gene is cut by Cas9.

Having established our initial approach to tagging, we applied it to gain insight into stress granules (SG), which are transient liquid-liquid phase separated (LLPS) RNA-protein complexes that form in response to environmental stress^12,13^. We hypothesized that by tagging a large number of SG-associated proteins with the fluorescent protein, mCherry, we could gain insight into the properties that drive some proteins to strongly accumulate within SGs. Toward this goal, a 193-member target-gRNA library against SG-associated proteins was designed and delivered to HEK293T cells (**Figure 1a**). Into the resulting mixture of cells, the remaining plasmids required for tagging were transfected in and puromycin-resistant cells were then selected. To determine the bias in tagging in the resulting library, the abundance of each gRNA within the population of cells before and after the procedure was determined (**Figure 1b**). As expected, the distribution of gRNAs in the initial population of cells before mCherry knock-in was even, with no gRNA dominating the pool. In contrast, after mCherry insertion and puromycin selection, a marked skew in the population was observed with 5 gRNAs occupying more than 60% of the pool, suggesting that only a handful of genes were efficiently tagged during our initial attempt and that further optimization was required.

### Optimizing insertion rates and drug selection steps

As numerous genes were poorly targeted in our initial library, we first sought to improve the efficiency of the process by optimizing the ratio of the transfected plasmids. Using a HEK293T cell line that stably expressed a single target-gRNA against the histone gene, HIST1H4C, 10ng or 100ng of each of the plasmids required for tagging (Cas9, donor plasmid, donor-gRNA) were delivered (**Figure 1c**). These studies revealed that a higher amount of the donor plasmid (100ng) and a lower amount of the donor-gRNA (10ng) led to a significant increase in the number of mCherry-positive cells, before the application of drug selection. Further optimization of the transfection conditions demonstrated an upper limit to the benefit of delivering more donor plasmid (500ng), improvements by adding additional Cas9 plasmid (100ng), and a decrease in insertion rates when the amount of donor-gRNA is further reduced relative to the amount of donor plasmid (**Figure 1c**). These results indicate that the amount of cleaved donor plasmid is a major factor determining the tagging efficiency and suggest that cutting the donor plasmid too early or too late via increased or decreased amounts of donor-gRNA, respectively is detrimental.

Given that the puromycin resistance marker is under the regulation of the endogenous gene that is tagged, we next hypothesized that weakly transcribed genes may not produce enough of the resistance protein to survive drug selection. To increase the level of puromycin resistance conferred upon tagging, a modified donor plasmid (P3) was constructed containing three tandem copies of the puromycin resistance gene downstream of the mCherry tag (**Figure 1d**). As a control, a donor plasmid with three copies of the puromycin resistance gene downstream of mCherry, but containing a stop codon after the first PuroR copy was also constructed (P3S). These modified donor plasmids along with the original donor plasmid with one copy of the PuroR gene (P1) were then transfected independently along with Cas9 and the donor-gRNA into HEK293T cells containing a single target-gRNA against one of three target genes, HIST1H4C, CCT7, or NUP93. Minimal difference in the number of mCherry-positive cells before puromycin selection was observed among the different donors (**Figure 1e**). In contrast, the number of viable cells dramatically increased after puromycin selection using the P3 donor plasmid which expresses the three copies of the puromycin resistance gene (**Figure 1f**). These results suggest that in our initial approach, many properly tagged cells were likely lost due to insufficient expression of the drug selection marker.

Upon applying the improved transfection conditions and optimized P3 donor plasmid design to the initial 193-member SG target-gRNA library, a marked improvement in the number of tagged genes was observed. Of the genes that were tagged in the pool, 29 genes showed an abundance >1% following puromycin selection as compared to only 8 genes meeting this threshold in the initial approach (**Figure 1b**). In addition, a strong correlation (_ρ_=0.78) among independent biological replicates was also found, suggesting that the tagging was efficient and reproducible (**Supplementary** Figure 6). We designated this optimized pipeline for generating pools of tagged cells HITAG.

### Application of HITAG to stress granule factors

To ensure optimal tagging, our initial library of 193 SG targets had been selected such that they all shared the same reading frame upon being cut by Cas9. To tag the remaining proteins previously associated with SGs, two additional target-gRNA libraries of 190 and 205 members were created and used to generate additional pools of mCherry-tagged cells via the HITAG approach.

To characterize the fidelity of HITAG, the junctions between the various target genes and the inserted mCherry tag (junction reads) were selectively enriched via nested PCR. From these analyses, 244 of the 588 genes that were targeted across the three reading frame libraries were tagged and survived drug selection (**Supplementary** Figure 7). The number of junction reads associated with each tagged gene was then compared to its respective target-gRNA abundance within the same pool of cells. As anticipated, a good correlation between these two metrics was observed (_ρ_ = 0.84), as would be expected if the target-gRNA within each cell was driving the tagging events. In addition, these data point to target-gRNA abundance being an easy-to-quantify surrogate for which genes are tagged within the library (**Figure 2a**).

**Figure 2.**
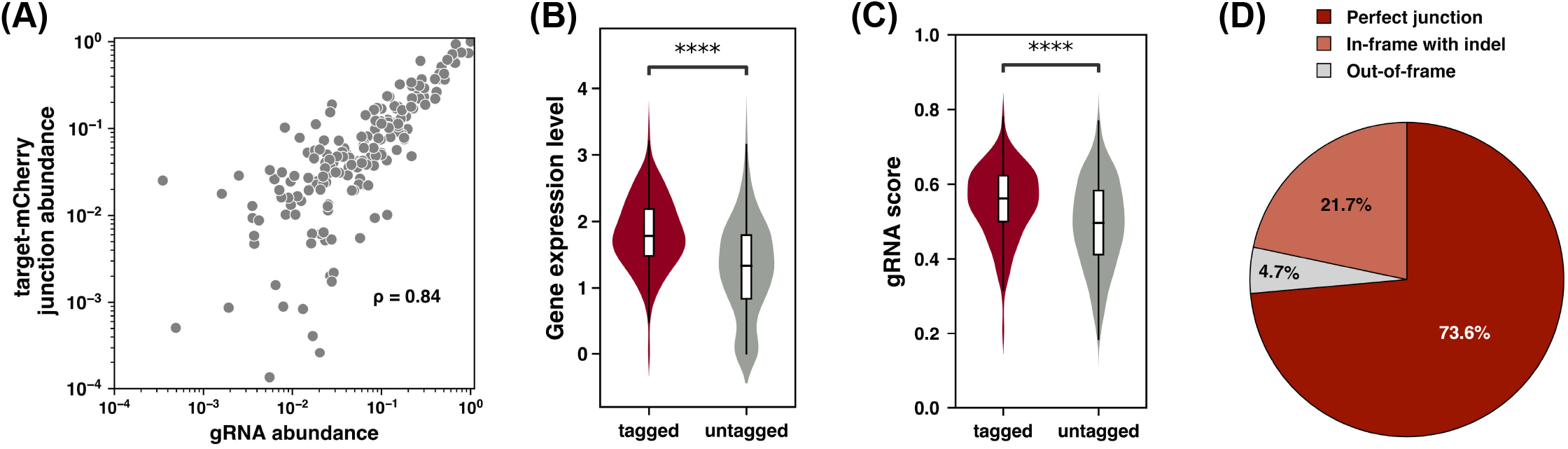
Characterization and validation of stress granule library generated with HITAG. (a) Correlation between the read counts from each gRNA within the pool of HITAG-modified cells compared with the target-mCherry tag junction reads derived from the same pool of cells. The abundance of each target-gRNA and junction reads within the pool of cells was internally normalized to be between 0 to 1 before performing the comparison. IZ represents the Spearman correlation coefficient between the two sets of data. Comparison of how (b) RNA expression level, represented as log (FPKM +1) or the (c) gRNA activity score of each target-gRNAs influences if a gene is tagged or not within the pool of tagged HEK293T cells (Mann-Whitney-Wilcoxon test, **** = p < 10^-4^). (d) Distribution of repair outcomes summed across all targets upon performing HITAG in HEK293T cells. Perfect junction indicates precise annealing of the endogenous locus to the donor plasmid without a loss or addition of bases after Cas9 cutting.

While HITAG was able to target a large proportion of the genes within the library, many failed to be tagged. To investigate the factors influencing these results, the characteristics of tagged and untagged genes were interrogated. Given our previous findings regarding the expression of the puromycin resistance marker limiting the efficiency of selection, we first examined if gene expression influenced tagging efficiency. On average tagged genes showed a higher level of expression as compared to untagged genes (**Figure 2b**). Despite the overall importance of gene expression, several genes showed high expression but remained poorly targeted. To reduce the number of amino acids lost from the C-terminus of our proteins of interest, gRNAs were selected based on their proximity to the stop codon. We hypothesized that a portion of the variation in tagging may be driven by inefficient target-gRNAs that poorly cut the endogenous target site. To investigate this hypothesis, we obtained a predicted activity score for each gRNA using CRISPick and found that tagged genes on average had target-gRNAs with significantly higher predicted activity^14^ (**Figure 2c**).

A concern when using NHEJ to insert DNA sequences into endogenous loci is that it can lead to error-prone repair with additional bases being lost from both the target gene and donor construct. When analyzing the sequenced junctions from the pool of mCherry-tagged cells, 73.6% of all junctions were precise fusions between the endogenous locus and the donor plasmid, in line with previous studies ^9,15^ (**Figure 2d**). Of the remaining junctions, 21.7% retained the proper frame between the gene of interest and the mCherry tag but showed a loss or gain of bases, with a strong bias towards smaller 1-2 amino acid deletions and insertions within the observed repair products (**Figure 2d** and **Supplementary** Figure 8). The remaining 4.7% of the repaired junctions were out-of-frame products, presumed to have arisen from cells that have both a properly tagged allele conferring drug resistance and an out-of-frame allele.

To investigate whether the HITAG approach can be applied to other cell lines, a set of 205 stress granule-associated genes were tagged with mCherry using the human colorectal carcinoma cell line, HCT116 (**Supplementary** Figure 9). The features of tagging such as the efficiency, specificity, overall repair profile, and biases in tagging were similar between HCT116 and HEK293T cells, in line with HITAG being a generalizable method for tagging genes in high-throughput.

### Characterization of tagged proteins and their association with stress granules

To examine the behavior of the mCherry-tagged proteins within our mixed pool, single cells were isolated, grown clonally, and the gRNA inside each was sequenced (for details see materials and methods). Within the 806 clonal lines obtained, 167 unique proteins were mCherry tagged, with each protein being represented by a median of 3 clonal isolates (**Supplementary** Figure 10). To quantify the rates of improper tagging, proteins that were tagged within at least 10 independent isolates and showed distinct subcellular localization (e.g. nuclear localization) were examined (**Supplementary** Figure 11a). On average ∼95% of the clones for a given protein showed the expected pattern of mCherry localization. These results were further confirmed for two targets, BCALF1 and HNRNPA2B1, by using PCR to amplify the junction between the target gene and the mCherry tag (**Supplementary** Figure 12b).

To probe if the mCherry tag alters protein localization, the subcellular distribution of the tagged proteins (**Supplementary** Figure 12) was compared with the annotations listed within the Human Protein Atlas^16^. Of the 155 proteins with localization data in the Human Protein Atlas, 141 agreed with our findings. For the 14 proteins with conflicting results, 6 had annotations in other databases^7,17^, with 5 of these agreeing with our findings^7^. As a whole, these data suggest that HITAG generates a high percentage of cells where the label is inserted as designed and that the C-terminal fusion of mCherry to proteins of interest rarely perturbs their localization.

While there are hundreds of proteins that have been found in SGs, what drives their accumulation and why some proteins are more efficiently recruited to SGs is under active investigation. To probe these questions, all 167 clonal lines were treated with 0.5 mM sodium arsenite for 1 hour to induce SG formation. Stress granules were then visualized by staining for the canonical SG marker G3BP1 (**Figure 3a**). Of the 167 tagged proteins, only 23 showed the formation of distinct foci that overlapped with the G3BP1 marker (strong accumulation), as determined by fluorescence microscopy (**Figure 3b**). To determine if the mCherry label might be affecting protein dynamics, 9 proteins that showed either strong or weak accumulation (no G3BP1 overlapping foci) within SGs were stained with antibodies against the target protein in wild-type cells and the results compared to our findings (**Supplementary** Figure 13). Comparisons between the staining patterns showed that the mCherry tag had a minimal effect on altering a given protein’s interaction with SGs.

**Figure 3.**
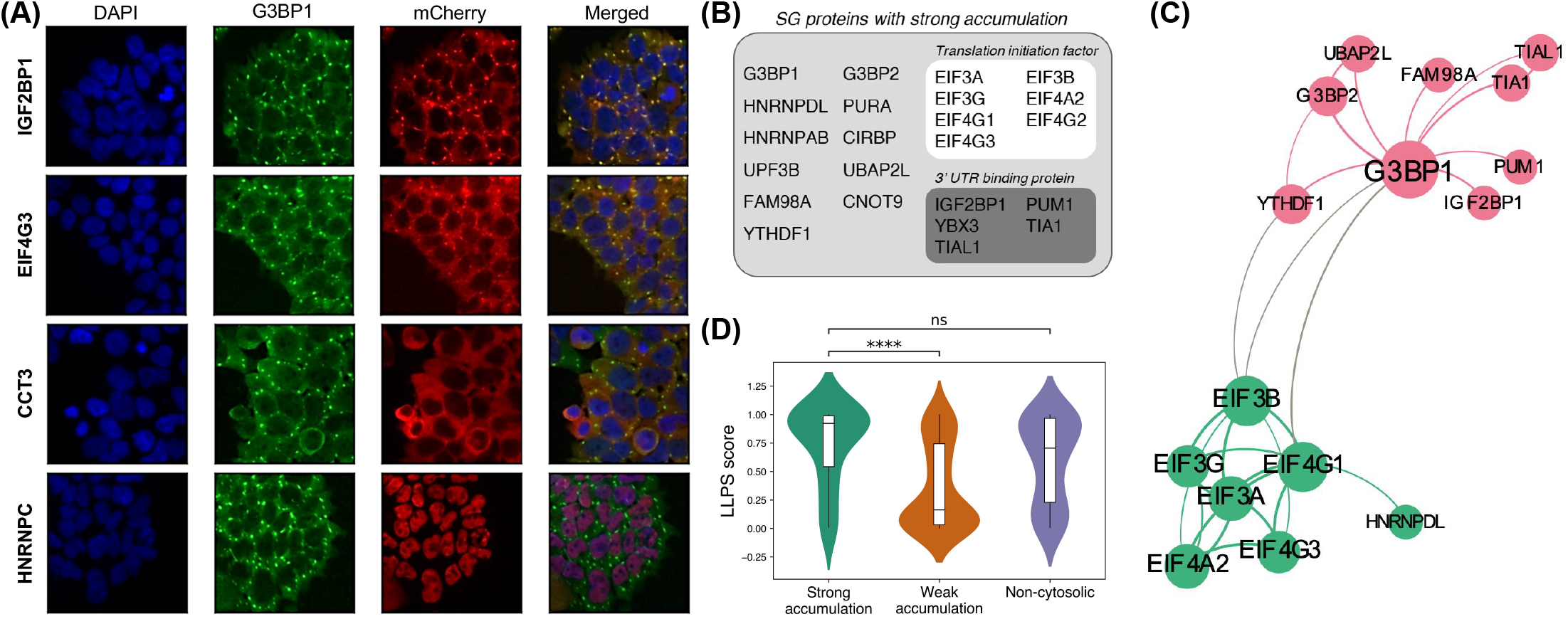
Using HITAG to understand the properties of proteins that accumulate within SGs. (a) Staining of proteins after inducing SG formation with the treatment with 0.5 mM of NaAs_2_O_3_ for 1 hour. (Blue:DAPI, Green: anti-G3BP1, Red: anti-mCherry). (b) List of proteins found to strongly accumulate within SG as determined by their overlap with the canonical SG marker, G3BP1. (c) Network depicting the interactions among proteins which show robust accumulation within SGs. (d) Predicted ability to LLPS as determined by LLPSDB. As all proteins that showed strong accumulation were predominantly localized to the cytoplasm for comparison cytoplasmic proteins were divided into those that strongly or weakly accumulate into SGs. As an additional group, we included all other proteins that showed weak accumulation but did not primarily localize to the cytoplasm (non-cytosolic) (Mann-Whitney-Wilcoxon test, ns = not significant; **** = p < 10^-4^).

In examining the 23 proteins with strong accumulation, several distinctive features immediately became apparent. Among the hits, all showed predominantly cytoplasmic localization and had associated RNA binding activity. In addition, the majority could be clustered by protein-protein interactions into two groups centered around either EIF4G1 or G3BP1 (**Figure 3c**).

To further characterize the nature of proteins that showed strong versus weak accumulation each of the proteins was compared on a variety of features such as protein length, size of intrinsically disordered regions, abundance, charge, and liquid-liquid phase separation (LLPS) propensity (**Figure 3d** and **Supplementary** Figure 14). Somewhat surprisingly, features such as protein length or the number of protein interaction partners showed no correlation with a protein’s strength of accumulation within SGs. Of the metrics examined, a propensity to LLPS showed the greatest difference between strong and weak accumulators, suggesting a critical role for multivalent intra and intermolecular interactions in driving these proteins’ behavior within SGs. These findings are in line with the fact that during times of translation inhibition, SGs form by stalled preinitiation complexes, containing EIF4G1, and their associated RNAs which undergo condensation upon interaction with key nucleating RBPs such as G3BP1^12^. They also highlight the known role of protein-RNA and protein-protein interactions in driving SG formation^12,13^.

As a whole, these results demonstrate that while many proteins may associate with SG, only a relative few have the required features to enable their robust accumulation. In agreement with this observation, several nuclear RNA-binding proteins were found to have high LLPS scores but poor accumulation within SG in our dataset. Yet, previous reports have shown that during times of stress, some of these proteins can become localized to the cytoplasm, fulfilling the critical requirement we highlight, which then enables them to readily accumulate within SGs^18,19^.

## Discussion

Here we report the development of High-throughput Insertion of Tags Across the Genome (HITAG), a fast and accessible method for generating libraries of endogenously tagged proteins. Although several methods exist for performing targeted knock-ins into the mammalian genome, most allow only a single protein at a time to be targeted, are inefficient, or place tags in between coding exons which can disrupt protein folding and function^7,8,20–23^. By increasing the throughput of performing C-terminal tagging, HITAG will help facilitate the scalable interrogation of protein function and dynamics. In future developments, the coupling of HITAG with single-cell approaches such as pooled optical screens will remove the need to isolate clonal populations post-tagging and provide a further boost to discovery throughput^24^.

Although all the proteins that were fused to mCherry in this study have been previously annotated as stress granule-associated, only 23 showed fluorescent puncta that overlapped with the canonical SG marker G3BP1 upon subjecting the cells to sodium arsenite treatment^25,26^. While several factors were found to be associated with strong accumulation within G3BP1 foci, cytosolic localization, the ability to interact with RNA, and a propensity to phase separate were found to be among the most critical. In future studies, the generated lines may be used in combination with live cell and single molecule imaging modalities to better understand the kinetics and distribution of each tagged protein within SGs to gain further insight into granule assembly, disassembly, and heterogeneity.

While HITAG can generate a diverse library of endogenously tagged proteins, several areas can be further explored to improve the quality of the generated libraries. First, the correlation between gene expression and tagging efficiency, in combination with the observation that knocking in multiple puromycin resistance genes reduces our targeting bias, suggests that lowly expressed genes are being lost due to insufficient levels of drug marker expression. To overcome this limitation, additional copies of the puromycin resistance marker, beyond the three currently being used, should be considered. In addition, efforts to identify more efficient drug markers via the mining of metagenomes or performing laboratory evolution on existing markers are additional options that can be considered^27,28^. Second, the finding that predicted gRNA activity influences the rate of gene tagging suggests that Cas9 variants with reduced PAM requirements may increase the efficiency of targeting by allowing greater flexibility in selecting which gRNA to use while still enabling insertions to occur near the C-terminus of the target genes^29,30^. In addition, although a perfect junction without any loss or insertion of amino acids is the most common repair outcome observed during the use of HITAG, there are still ∼22% of junctions that show a loss or gain in nucleotides. This raises the possibility that one may be able to direct Cas9 to cut downstream of the stop codon and rely on error-prone repair to process away the stop codon and place the tag in-frame of the protein of interest. If this alternative approach was found to be efficient, it would increase the number of gRNA available for use and may also reduce the number of amino acids that are lost during tagging, which at present is on average two.

## Acknowledgments

This work was supported by a Career Awards for Medical Scientists from the Burroughs Wellcome Fund and NIH grant 1R21HG011855-01 to A.C. Flow cytometry experiments were performed in CCTI supported by core-supporting NIH grant S10RR027050. Cell sorting experiments were performed in CSCI Flow Cytometry core facility at Columbia University Irving Medical Center under the leadership of Michael Kissner.

## Contributions

J.K. and A.C. designed experiments, assisted with data interpretation, and directed the research program. B.K.S. performed the initial feasibility studies. J.K. optimized the HITAG methodology including the development of the triple puromycin marker. J.K. and J.S. isolated the clonal population. J.K., J.S., and S.C. conducted the NGS sample preparation. J.K., J.S., A.F.K., L.Z., and S.C. analyzed the resulting NGS data. J.K., L.Z., and S.C. performed the image analysis and interpreted the results of the localization data. J.K. and A.C. wrote the manuscript with input from all authors.

## Conflicts of interest

A.C. has several issued and provisional patents managed by Harvard University and Columbia University on CRISPR-based tools and methods. In addition, AC, JK, and BKS have filed patents related to HITAG with Columbia University.

## Data availability

All sequencing data were uploaded to SRA under BioProject accession number: PRJNA895413. The script for the junction read analysis is available at https://github.com/afkratz/HITAG-analysis-code. Additional scripts can be obtained upon written request.

## Materials and Methods

### Plasmid construction

To construct the gRNA expression plasmids, pSB700-blasto (Addgene #167904) was used for SpCas9-specific gRNA expression and a modified pSB700-vector containing a SaCas9 compatible gRNA scaffold with a zeocin-resistance gene was used for SaCas9-specific gRNA expression. Vectors containing gRNAs were cloned by Golden Gate using Esp3I as described previously^31^. pCAS plasmid was constructed from a previous dual-Cas9 plasmid (Addgene #107320) by replacing the 3×HA sequence with a P2A sequence using Gibson assembly. pDNR was constructed from pCRISPaint-TagGFP2-PuroR (Addgene #80970) using Gibson assembly. To construct the modified P3 donor with additional copies of the puromycin resistance gene two puromycin resistance genes were PCR amplified with primers designed to add a T2A sequence on each end. These fragments were then assembled into a version of pDNR that was digested with ZraI using Gibson assembly. All plasmids were validated by Sanger sequencing and will be made available via Addgene. The sequence of the P3 donor, pCAS, the target-gRNA, and donor-gRNA vectors are provided in **Supplementary Table 2**.

### Target-gRNA design

To design target-gRNAs the CRISPick tool from the Broad was used with settings Human GRCh38, CRISPRko, and SpyoCas9^14^. Guide RNAs with Esp3I restriction sites inside of them, a polyT stretch longer than 4 base pairs, or more than one exact match in the human genome were excluded from use even if they were the gRNA closest to the stop codon. Frame number was categorized as the number of bases required to complete the cut codon after cleavage.

### Construction of stress granule library

A list of stress granule-associated genes was collected from the existing literature^25,26^. These methods identified SG proteins based on the biotinylation of proteins in close spatial proximity to core SG components or affinity purification of SG followed by mass spectrometry. Using this list a set of target-gRNAs against each gene was generated with any target-gRNAs that would result in a loss of more than 6 amino acids from the C-terminus of a protein being removed. The resultant list of target-gRNAs was then sorted into three libraries based on the frame (**Supplementary Table 1**).

Three gRNA libraries with different frames were synthesized as oligo pools from Agilent. Each gRNA library was then PCR amplified from the initial oligo pool and cloned into the modified pSB700 vector using Golden Gate assembly as previously described^31^.

### Construction of HEK293T cells containing a library of target-gRNAs

HEK293T cells were cultured in Dulbecco’s Modified Eagle Medium (DMEM) + 10% FBS + 1% penicillin/streptomycin and incubated at 37°C and 5.0% CO_2_. On day 1, HEK293T cells were seeded at ∼3.5×10^6^ cells per well in 6-well plates for lentivirus production. On day 2, a mixture of 600ng psPAX2, 150ng pMD2.G, and 450ng of the target-gRNA plasmid was transfected into each well using lipofectamine 2000 according to the manufacturer’s instructions. On day 3, the media was changed. Lentivirus was harvested by collecting the supernatant after centrifugation (500g for 5 minutes) of the media on day 4 and day 5. The lentivirus stocks were stored at –80°C in 1ml aliquots. To generate HEK293T cells with on average a single target-gRNA integrated into their genome viral stocks were tested for infectivity and cells were infected at an MOI of ∼0.1.

### Quantifying the efficiency of tagging

When optimizing the tagging approach, cells were plated at a density of 5-7×10^4^ cells per well in a 24-well plate with a media volume of 600µl. After 24 hours transfections were performed according to the manufacturer’s instructions with 100ng pCAS, 100ng pDNR-gRNA, and 200ng pDNR being delivered. In cases where the amount of each vector being delivered was changed an inert “stuffer” plasmid, pBR322, was delivered to ensure that the same total amount of plasmid DNA was delivered across all conditions. Media was changed on the next day of transfection.

On day 6 after transfection, the transfected population was split to poly-D-lysine (PDL)-treated 96 well plates at a density of 1-2×10^4^ cells per well. The following day cells were fixed with 4% paraformaldehyde for 5 minutes. The cells were then washed with PBS, stained with DAPI for 5 minutes, and washed twice with PBS. The cells were imaged using the ImageXpress® Pico Automated Cell Imaging Systems. Tagging efficiency was determined by the number of mCherry-positive cells showing the proper localization over the total number of cells as determined by DAPI staining of nuclei.

### Creating HITAG libraries

To initiate the HITAG procedure target-gRNA containing cell libraries were seeded into 6-well plates at 2.5-5.0×10^5^ cells per 6-well. One 6-well plate was used per library. On the next day, HITAG plasmids were transfected with 5 times more plasmid and transfection reagents used for the 24-well plate condition. Cells were split 1:4 in media with 0.5µg/ml puromycin 7 days after the transfection. The selection stopped until the next day when the negative control cells died out. Selected tagged cells were expanded by changing media.

A stress granule library of HCT116 cells was generated by going through the same procedure as outlined for HEK293T cells except for the concentration of drug (blastidicine, 5.0µg/ml; puromycin, 2.5µg/ml), transfection reagent (FugeneHD), and the scale at which transfections were performed (4 T75 flasks) was varied.

### gRNA analysis

1-2×10^6^ cells were harvested with trypsin and washed once with PBS. The cell pellet was resuspended in 500µl Lucigen DNA QuickExtract reagent and incubated at 65°C with shaking at 750rpm for 15 minutes. After brief centrifugation, the sample was incubated at 95°C with shaking at 750rpm for 5 minutes. gRNA regions were PCR amplified using the following conditions: 98°C 45s, [98°C 15s; 56°C 15s; 72°C 20s] x n, 72°C 2min, 4°C hold, where n is the cycle number before the PCR reaction saturated (usually between 20-25 cycles). The second round PCR was performed to add the Illumina indices: 98°C 45s, [98°C 15s; 56°C 15s; 72°C 20s]x8, 72°C 2 min, hold at 4°C. The PCR products were then run on a 2% agarose gel and the band of interest was purified. Samples were then sequenced on an Illumina NextSeq 500. The resulting reads were then analyzed by either aligning them using Bowtie2 or the MAGeCK pipeline^32^.

### High-throughput isolation and analysis of target-mCherry junctions

Genomic DNA was extracted from 1-2×10^6^ cells using the Qiagen DNA extraction kit (#69504). Enzymatic DNA fragmentation was performed using the NEBNext® Ultra™ II FS DNA Library Prep Kit (E7805S). 2µg of genomic DNA (500ng per reaction) was treated with 5 minutes of enzymatic fragmentation. All subsequent steps were performed as instructed by the manufacturer. The DNA fragments containing the mCherry sequence were amplified through a nested PCR approach. For the first round PCR, 500ng of fragmented and adapter-ligated DNA was amplified (50ng per reaction) using a primer binding on the reverse strand of the mCherry sequence (see **Supplementary Table 3** for primer sequences) and another primer binding to the p7 adaptor using the following PCR condition: 98°C 45s, [98°C 15s, 65°C 15s, 72°C 90s] x20, 72°C 5min, hold 4°C. After the resulting sample was purified using SPRI beads aiming to capture all products greater than 100 bp in size. The second-round PCR reaction was then performed using 50 ng of the first-round PCR product (10ng per reaction) using PJ3 (**Supplementary Table 3**) and index primers under the following PCR condition: 98°C 45s, [98°C 15s, 65°C 15s, 72°C 90s] x8, 72°C 5min, hold 4°C. The final PCR products were then isolated using SPRI beads aiming to remove all products smaller than 200 base pairs.

To characterize the target gene-mCherry tag junctions, we began by constructing a database of all genomic regions adjacent to the target cut sites. NGS reads were then blasted locally twice, once against the database of target genomic regions, and once against the three linker sequences, which differ by a few bases to maintain the appropriate reading frame. Reads which featured at least 20 bases of alignment to genomic targets and at least 20 bases of alignment to linker sequences were analyzed by comparing the locations of the alignments to the expected location based on the gRNA cut site. Insertions were identified as sections of a read which did not align with either the genomic target or the linker sequence, while deletions appeared as a difference between the expected cut site and the observed cut site.

### Generation of clonal cell lines and identification of the gRNA in each well

The library of cells with mCherry-tagged SG-associated proteins was detached from a T75 flask and washed once with prechilled PBS. The cells were resuspended in Ca/Mg-free PBS + 1% FBS, filtered with a 50um mesh filter, and kept in ice. Before single-cell sorting, SYTOX Blue (1:1000, Thermo S34857) was added as a cell viability indicator. In preparation for single-cell sorting, 96-well plates were filled with 150µl of media and prewarmed to room temperature. Viable cells were sorted on the 96-well plates using Sony MA900 in the Single-Cell Mode. Cells were placed in the incubator and the media from all plates was changed as needed. Each well was confirmed visually to have one colony per well after 10 days.

PCR-ready genomic DNA was prepared by mixing ∼2×10^4^ cells in each well with 30µl Lucigen DNA QuickExtract. After incubating the plates for 15 minutes at 65°C followed by 10 minutes at 95°C, 1µl of the DNA extract was used for PCR to amplify the gRNA sequence. The same gRNA PCR conditions were used as described above except 35 cycles were used during the first round of PCR. Read counts of gRNAs for each well were then analyzed. For a gRNA to be identified in a given well it needed to be present at an abundance at least 3 times greater than the next most abundant gRNA in that same well.

### Immunofluorescence staining for stress granule formation

Sodium arsenite stress was applied by incubating cells with media containing 0.5mM NaAs2O3 for an hour. Cells were then washed once with PBS and immediately fixed with 4% paraformaldehyde for 5 minutes, washed twice with PBS + 0.1% TritonX-100 incubating for 10 minutes between each wash step, and blocked with Superblock^TM^ Blocking Buffer (Thermofisher #37581) with 0.1% TritonX-100 for 2 hours at room temperature. For primary antibody staining, cells were covered with 100µl Superblock with 0.1% TritonX-100 and primary antibodies (G3BP1, proteintech 13057-2-AP, 1:1000 dilution; mCherry, proteintech 68088-1, 1:1000 dilution) overnight at 4°C. After washing the cells with PBS + 0.1% TritonX-100 (15 minutes) and Superblock with 0.1% TritonX-100 (15 minutes), cells were incubated with secondary antibodies (Goat anti-mouse, Invitrogen A32727; Goat anti-rabbit, Invitrogen A32731, 1:1000 dilution) for an hour at room temperature. After one wash with PBS + 0.1% TritonX-100, cells were stained with 0.1 _μ_g/ml DAPI in PBS for 5 minutes followed by two PBS washes. At this point plates were either immediately imaged or covered with aluminum seals and stored at 4°C.

### Collection of protein features

Three published algorithms including PSPredictor, CatGranule, and Plaac were used to predict various scores related to phase separation^33–35^. The number of intrinsically disordered regions, number and fraction of charged residues, hydropathy, and net charge were calculated using CIDER^36^. All the scores were predicted with default parameters using the dominant isoform.

### Protein-protein interaction network analysis

The protein-protein interaction network was extracted from the STRING database, with the network type set as physical network and a minimum required interaction score of 0.400^37^. All of the text mining, experimental data, and databases were accepted as active interaction sources. Orphan genes, without interacting partners, are not included in the final network. K-means was used for clustering, and the cluster number was set to 2. Visualization is made by Gephi 0.9.2, with different colors indicating different gene groups and node size indicating node degree.

## Figure Legends

**Supplementary Figure 1.**
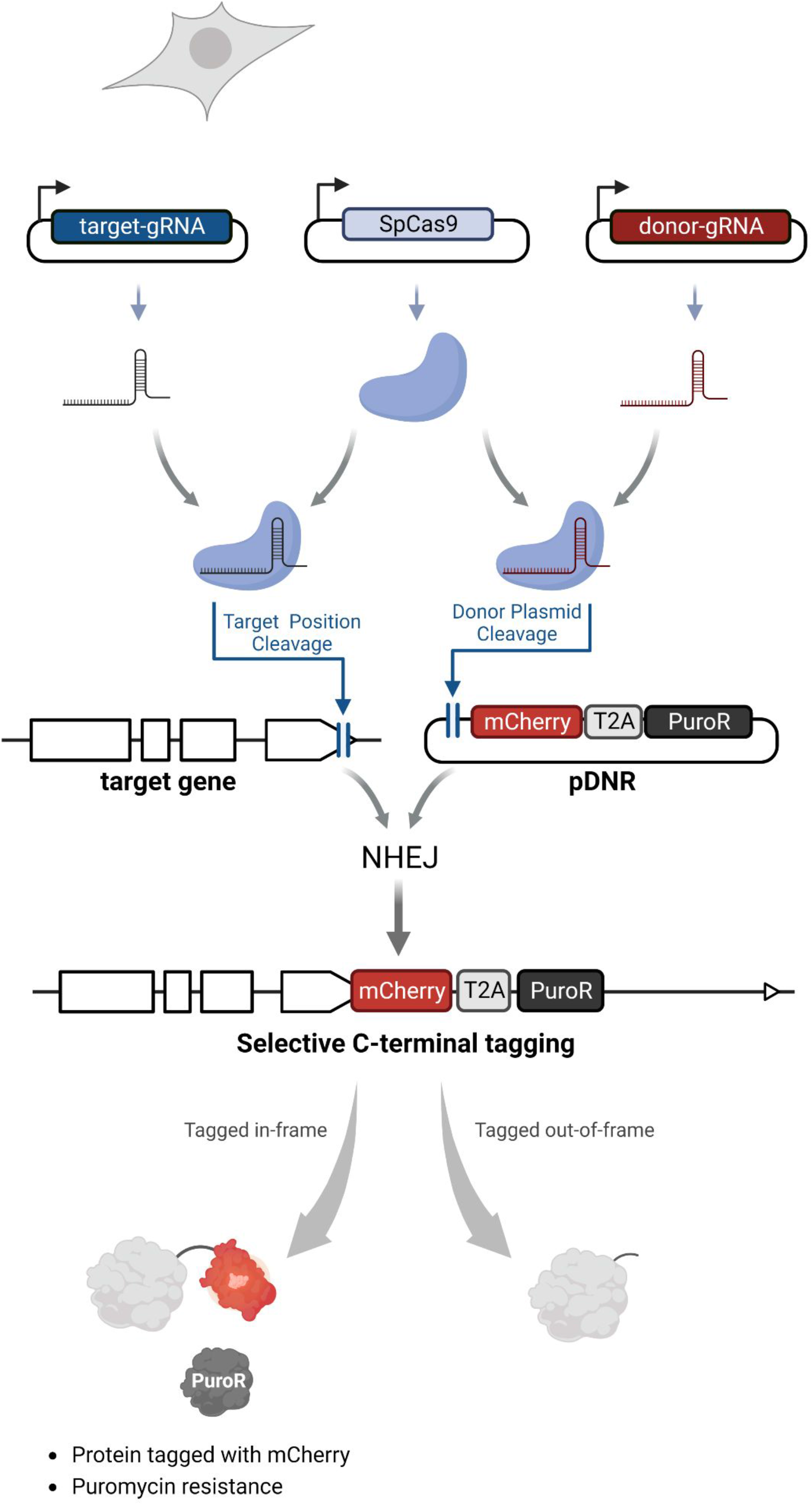
Schematic summary of CRISPaint approach for NHEJ-based gene tagging. Four different components – Cas9, target-gRNA, donor-gRNA, and donor plasmid – are transfected into cells and required for the CRISPaint approach. In CRISPaint, SpCas9 cleaves the target gene at its C-terminus and the donor plasmid (pDNR). Linearized pDNR can then become inserted into the genome through NHEJ. The cells which are properly tagged in-frame will express mCherry fused to the protein of interest along with the puromycin resistance gene (PuroR), which enables properly tagged cells to grow in the presence of puromycin.

**Supplementary Figure 2.**
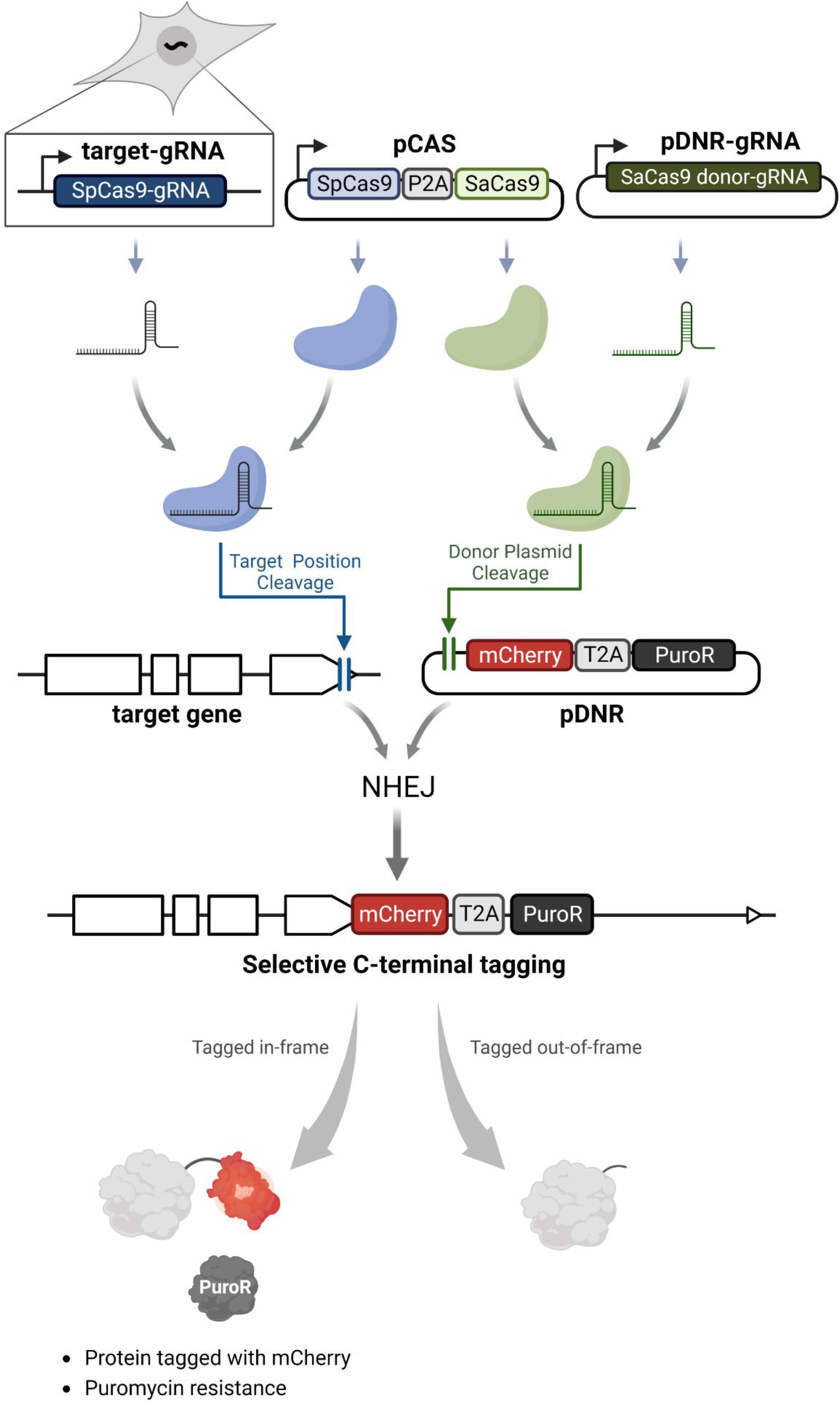
Schematic summary of HITAG approach for NHEJ-based high-throughput gene tagging. In the HITAG approach, the target-gRNA is designed to interact with SpCas9, while the donor-gRNA is designed to interact with SaCas9. The library of target-gRNAs against the various genes of interest is first integrated into the pool of cells at low infectivity such that each cell gets on average a single target-gRNA or none. The remaining CRISPR components are then transfected in. This then enables each cell in the population to tag a unique gene to which their target-gRNA is against. When tagging occurs in-frame the protein of interest gains the C-terminal tag and a drug resistance marker (e.g. puromycin resistance gene) is also expressed, enabling properly tagged cells to be readily isolated via a round of drug selection.

**Supplementary Figure 3.**
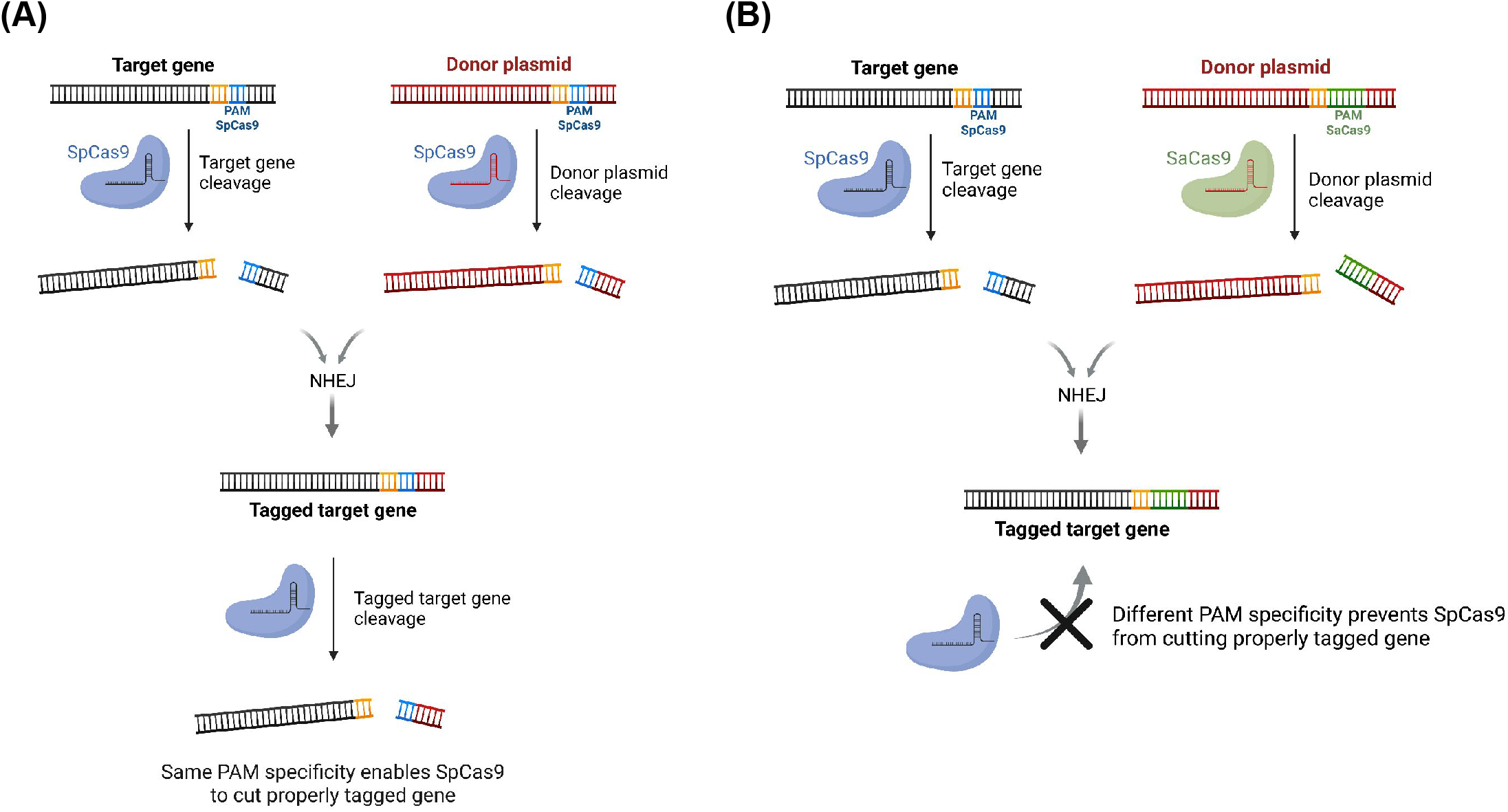
Schematic depicting how properly tagged genes can be recut when using only a single Cas9 protein. (a) When using only a single Cas9 protein, as is done in the original CRISPaint approach, the properly tagged product can be recut if the 3 PAM-proximal base pairs of the target gene are identical or similar to those of the donor plasmid. This occurs because the spacer sequence of the target-sgRNA with an accessible SpCas9-PAM site is regenerated in the final knock-in product. This recutting is then expected to inhibit the emergence of perfectly tagged products. (b) Recutting can be avoided when using two orthogonal Cas9 proteins, as is done in the HITAG approach. This occurs because even if the final product that forms has homology to the spacer sequence of the target-gRNA it will have a SaCas9-PAM site proximal to it, and thus not be an appropriate substrate for SpCas9 which has different PAM requirements.

**Supplementary Figure 4.**
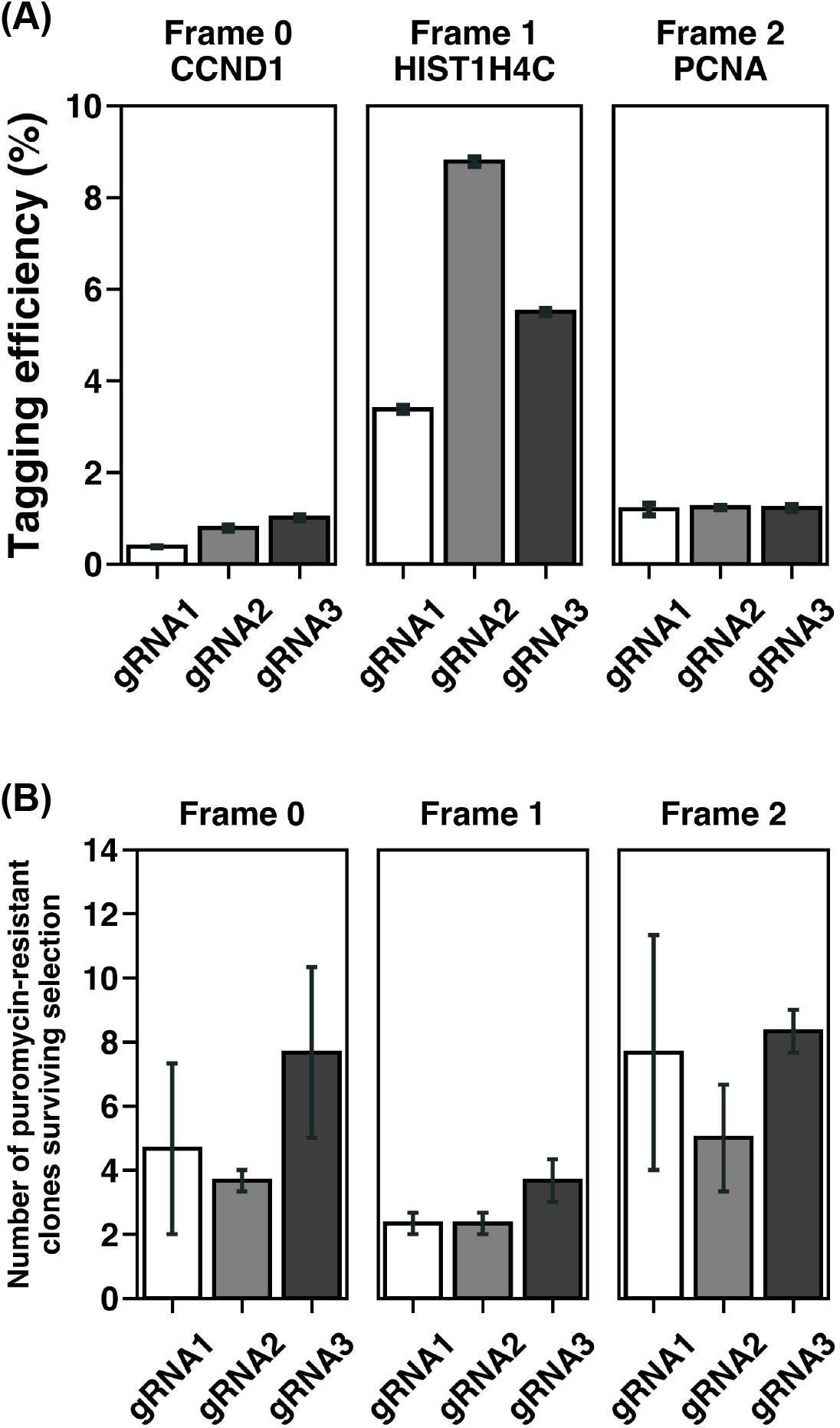
Identifying efficient and specific gRNAs against the donor plasmid. (a) The efficiency of three different donor plasmids targeting gRNAs (gRNA1-3) was assessed for the ability to add a C-terminal mCherry tag to either CCND1, HIST1H4C, or PCNA. To measure efficiency, the percentage of cells with the appropriate mCherry localization as determined by fluorescence microscopy was quantified before the application of drug selection. Frame refers to the reading frame of the donor plasmid when cut. As there are three possible reading frames (i.e 0, 1, and 2) that a given target gene could be cut in, 3 donor plasmids corresponding to each frame were prepared for each donor-gRNA being tested. This enables the system to then cut the target gene and donor plasmid in such a way that should they undergo a perfect fusion (without the addition or removal of bases) the tag and drug marker will be in the appropriate reading frame. (b) The same donor-gRNAs as in panel A were also examined for specificity by transfecting them in combination with SaCas9 and a mCherry-containing donor plasmid. As no target-gRNA is provided in this experiment, all drug-resistant clones that arise are due to either low levels of stochastic double-strand breaks or the donor-gRNA cutting an endogenous site in the genome to enable the donor plasmid to then insert itself. Based on these studies gRNA2 was determined to be the optimal donor-gRNA and for all subsequent studies was used. Three biological replicates (independent transfections) were performed for all conditions and error bars represent ± standard deviation.

**Supplementary Figure 5.**
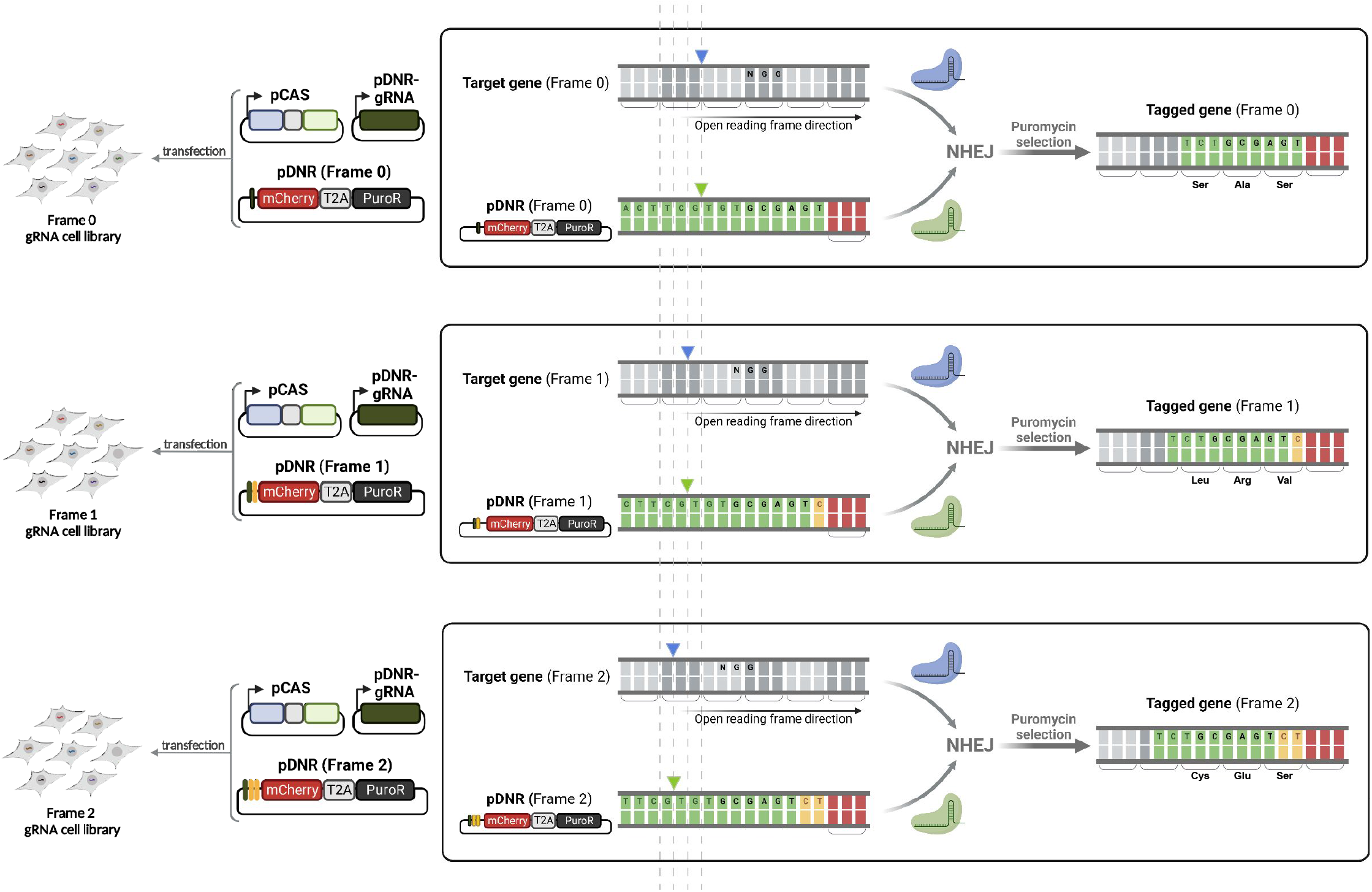
Schematic summary of the results of tagging depending on the “frame” in which the target gene is cleaved by SpCas9. Here, the frame number is defined as the number of nucleotides that must be added by the donor so that the target gene and the tag of interest are in-frame. In the original CRISPaint approach, three unique donor-gRNA are used against a single donor plasmid to generate each of the needed reading frames, with each donor-gRNA potentially having its own unique set of off-target within the genome. In contrast, HITAG employs three different donor plasmids that all work with a single donor-gRNA. By designing each donor plasmid to have either 0, 1, or 2 bases added it enables the same donor-gRNA, which is tested to ensure it has minimal off-target activity, to be used across all studies. For example, pDNR (Frame 0) is used when tagging a target gene that does not require additional nucleotides added for the tag to be in-frame with the cut gene.

**Supplementary Figure 6.**
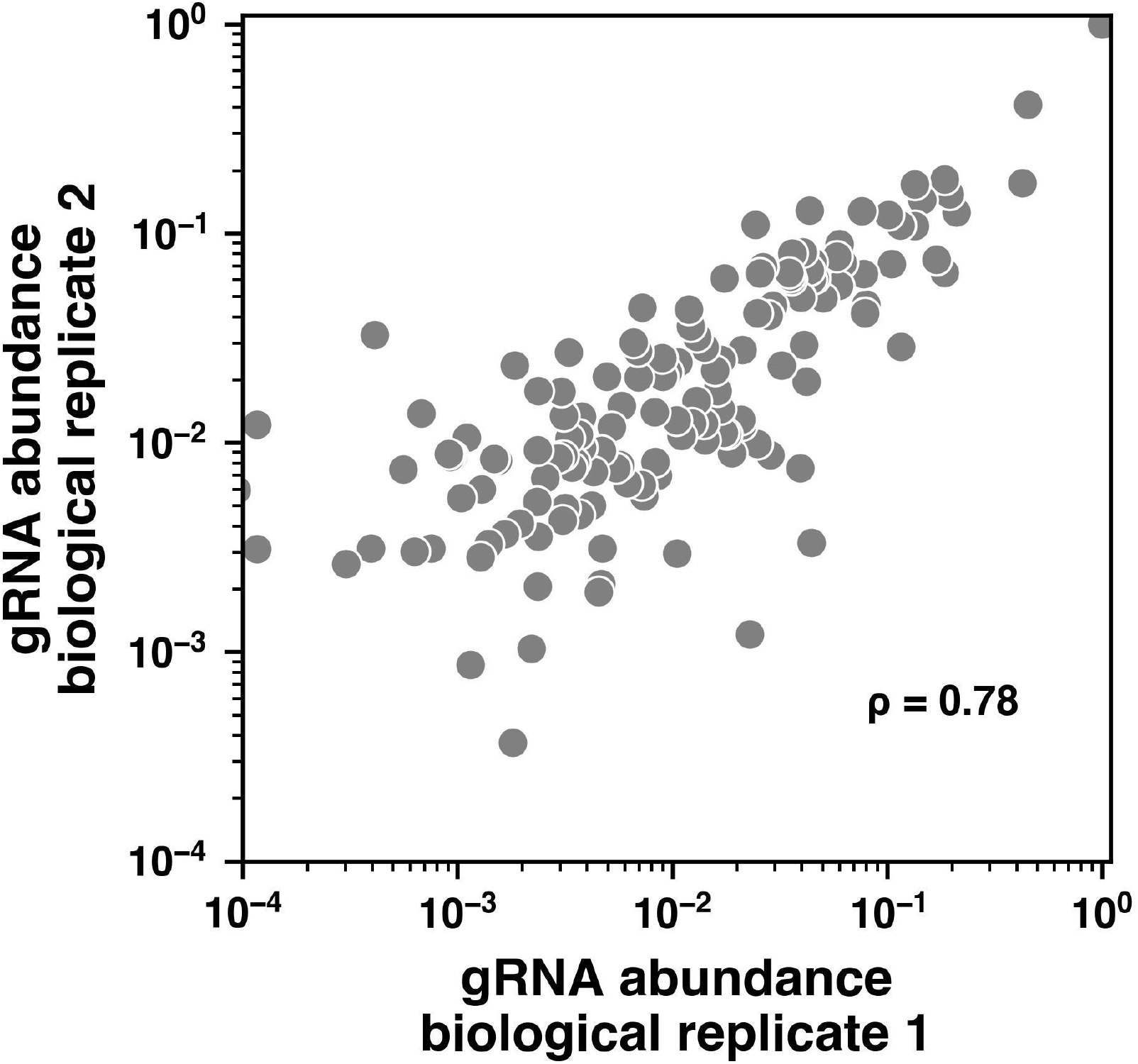
Comparing gRNA abundance between two independent rounds of HITAG on the same target-gRNA library. The abundance of each target-gRNA within the pool of cells was internally normalized between 0 to 1 for each replicate. IZ represents the Spearman correlation coefficient.

**Supplementary Figure 7.**
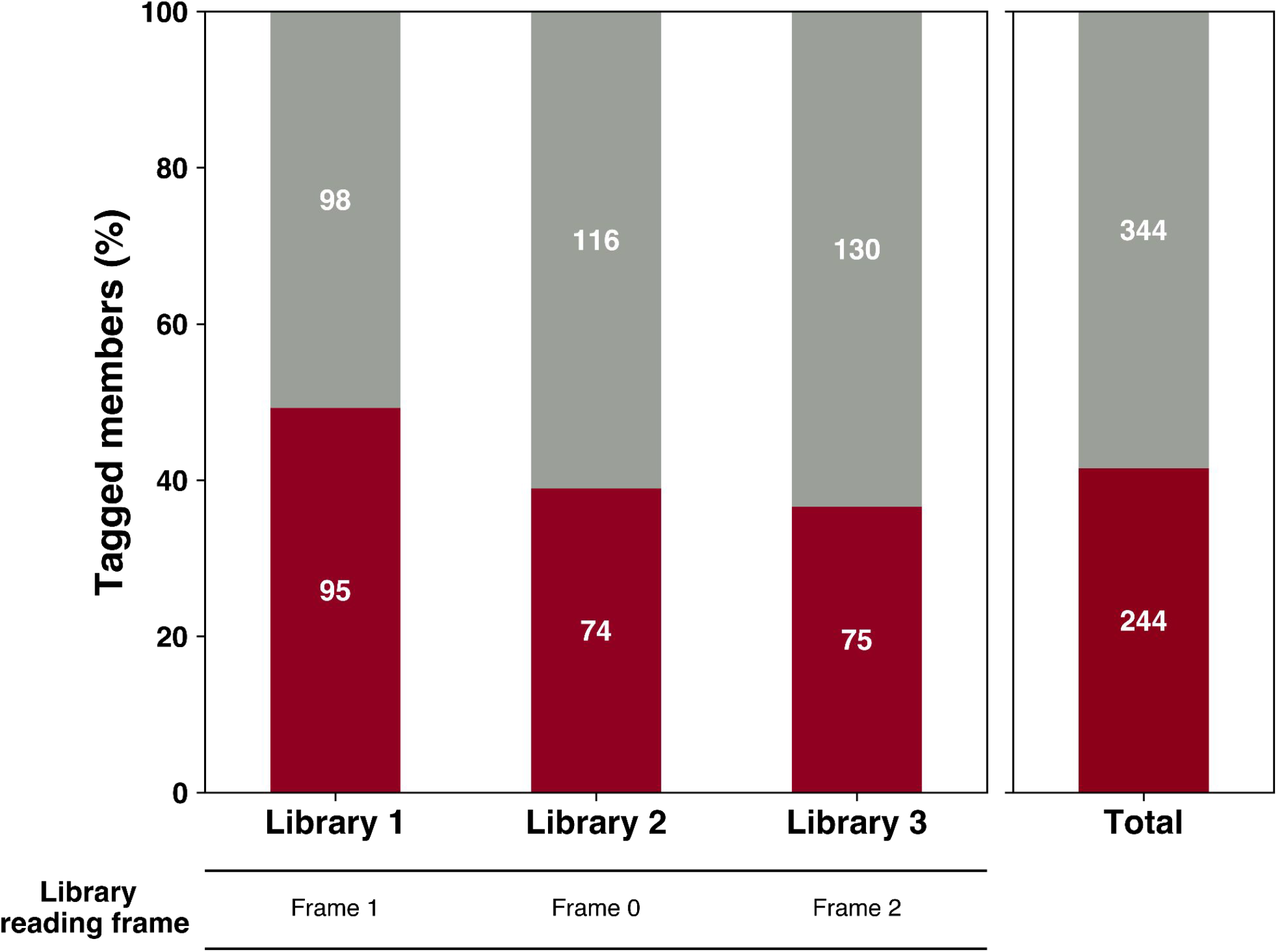
Percentage of tagged gene members within each reading frame and as a whole across the generated HITAG libraries. Red and grey boxes indicate the tagged group and the untagged group of genes, respectively.

**Supplementary Figure 8.**
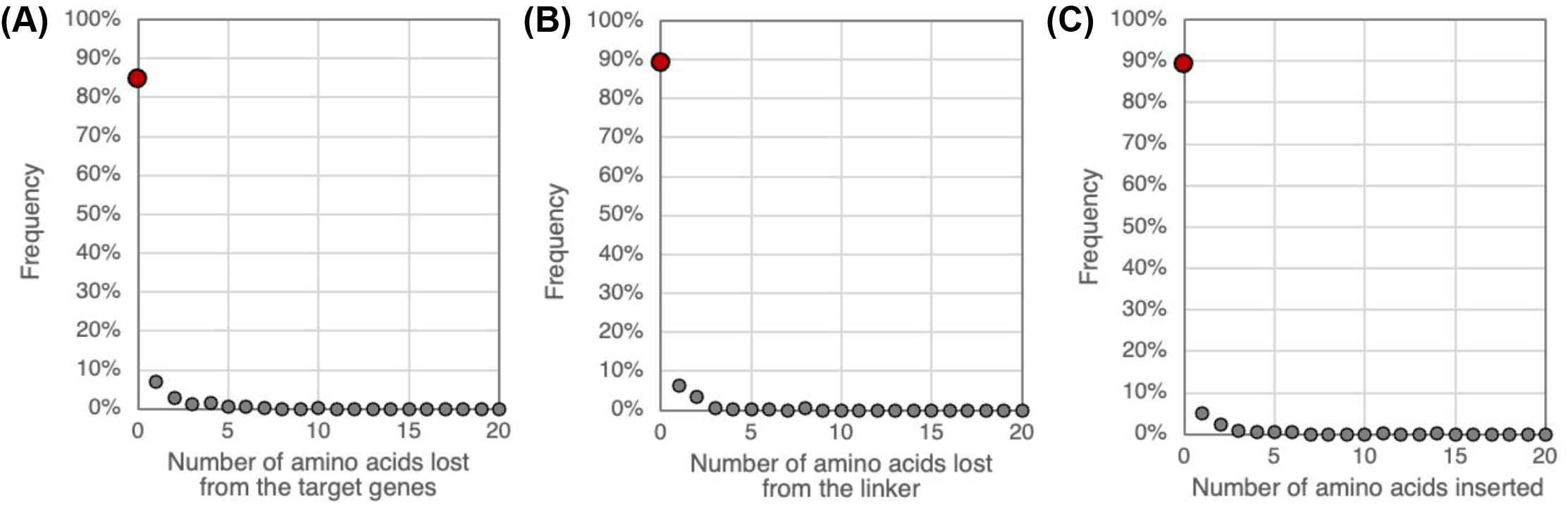
Characterization of the junction between the target gene and mCherry tag. Frequency of the number of amino acids lost from (a) the target genes or (b) the linker, which connects the tag to the endogenous target gene, respectively. (c) Frequency of the number of amino acids insertion. Insertion refers to the number of amino acids that are inserted between the target gene and the tag. Red dots indicate the junctions which show no deletion or insertion of additional amino acids.

**Supplementary Figure 9.**
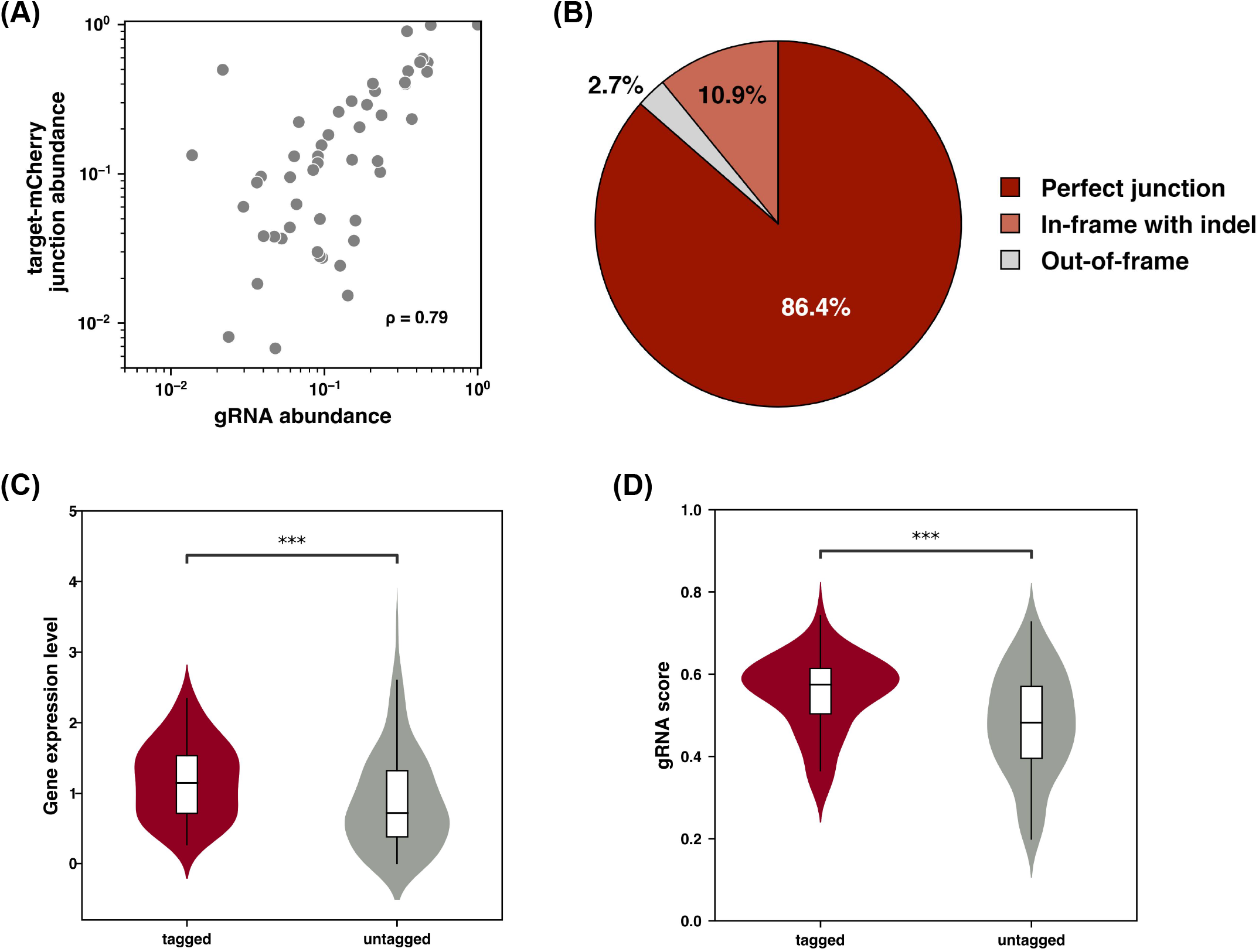
Result from applying HITAG to HCT116 cells. The SG target-gRNA library 3 (frame 2) was integrated into HCT116 cells and the subsequent pool of cells was taken through the remainder of the HITAG procedure to isolate a mixed population of mCherry-tagged cells. (a) Correlation between the normalized read counts from each gRNA within the pool of HITAG-modified HCT116 cells compared with the normalized target-mCherry tag junction reads derived from the same pool of tagged HCT116 cells. IZ represents the Spearman correlation coefficient between the two sets of data. (b) Distribution of repair outcomes summed across all targets upon performing HITAG in HCT116 cells. Perfect junction indicates precise annealing of the endogenous locus to the donor plasmid without a loss or addition of bases after Cas9 cutting. Comparison of how the (c) RNA expression level represented as log (FPKM +1) or the (d) On-target efficacy score of the target-gRNAs influence if a gene is tagged or not within the pool of tagged HCT116 cells (Mann-Whitney-Wilcoxon test; *** = p < 10^-3^).

**Supplementary Figure 10.**
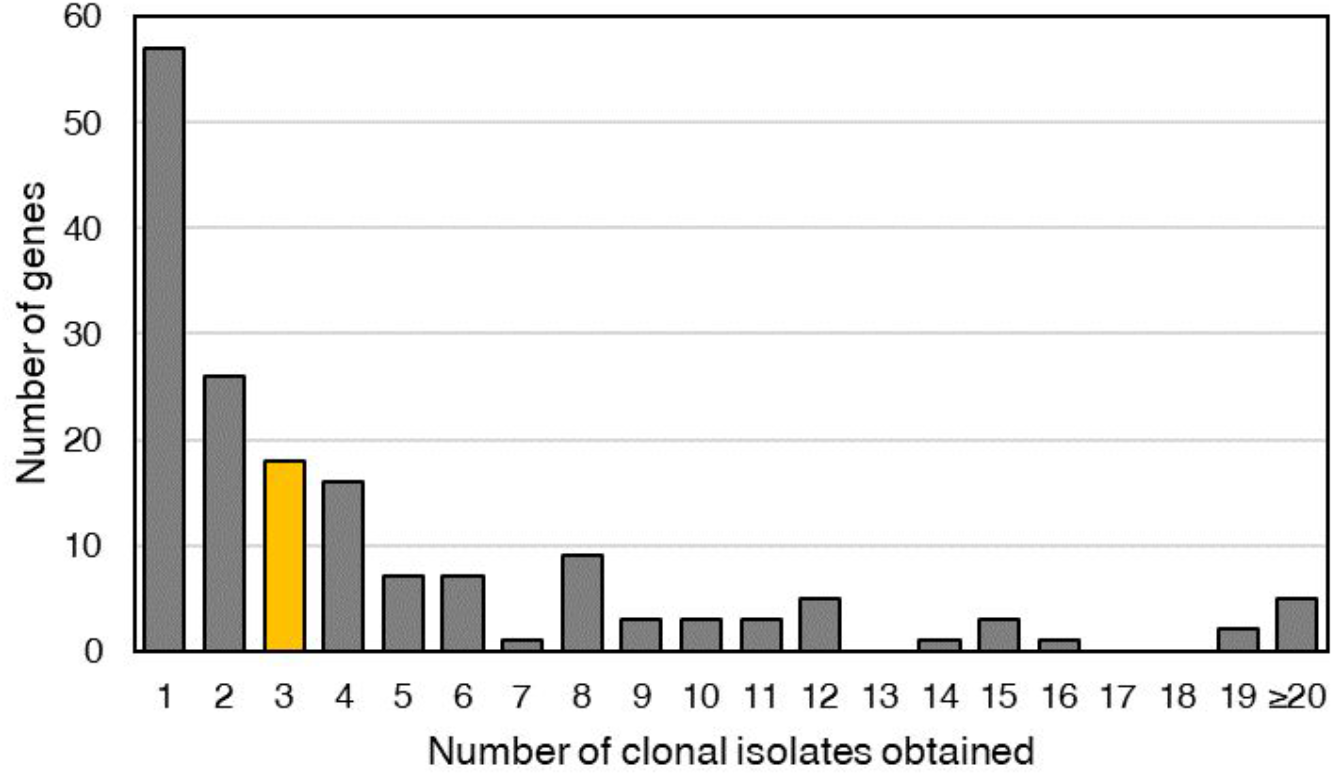
The number of times a given gene was tagged within the set of 806 clonal lines that were isolated after performing HITAG. The yellow bar represents the median number of clones obtained for a given targeted gene.

**Supplementary Figure 11.**
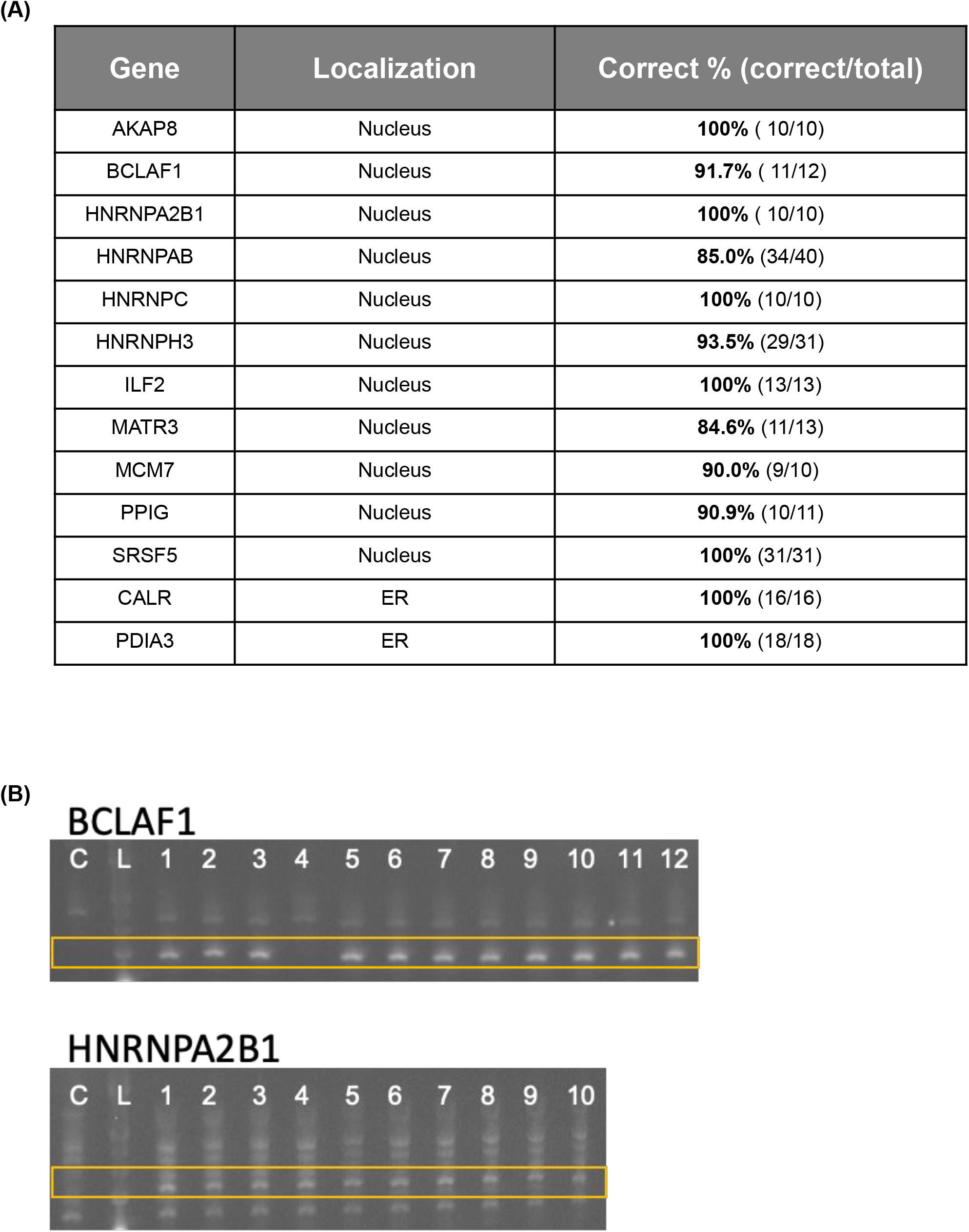
Examining the rates of off-target tagging by quantifying the consistency of results when multiple clones of the same tagged gene were obtained. (a) Proteins with either nuclear or ER localization which had 10 or more clones to examine from within the 806 clonal isolates were studied to determine the number of clones that showed the appropriate localization. (b) PCR validation using primers to show that microscopy-based results were concordant with targeted PCR directed at the junction between either HNRNPA2B1 or BCLAF1 and the mCherry tag. The yellow rectangle marks the location of the expected PCR band. C: control DNA from untargeted HEK293T cells. L: Ladder.

**Supplementary Figure 12.**
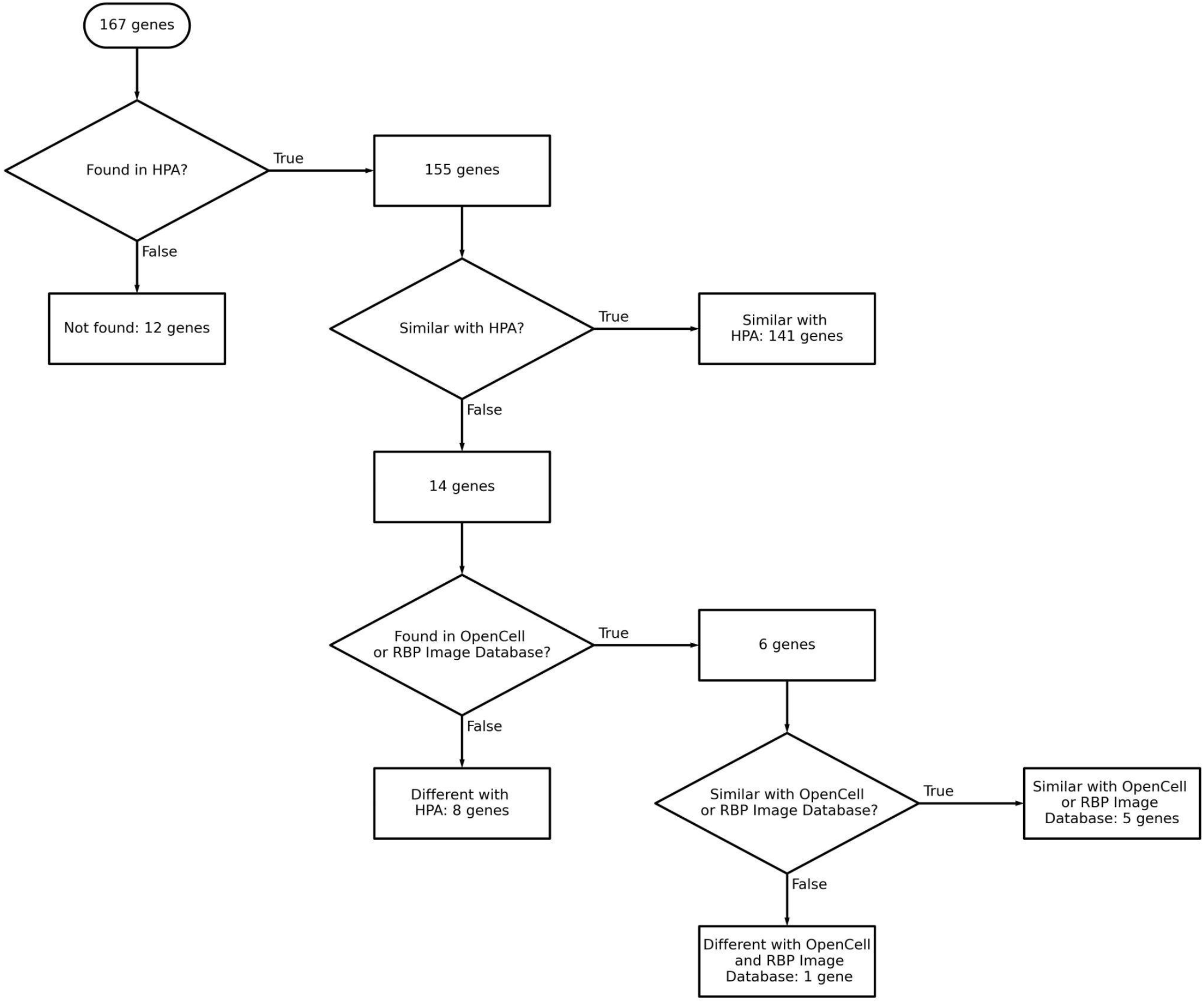
Decision tree used to compare localization information between this study and the Human Protein Atlas. Protein subcellular localization information was available for 155 out of 167 genes in Human Protein Atlas database. 141 out of 155 genes showed a similar subcellular localization within our data as described in Human Protein Atlas. Of the 14 genes that disagreed with the data from Human Protein Atlas, 6 genes were found in Opencell or the RBP Image Database. 5 out of 6 of these genes agreed with our subcellular localization findings.

**Supplementary Figure 13.**
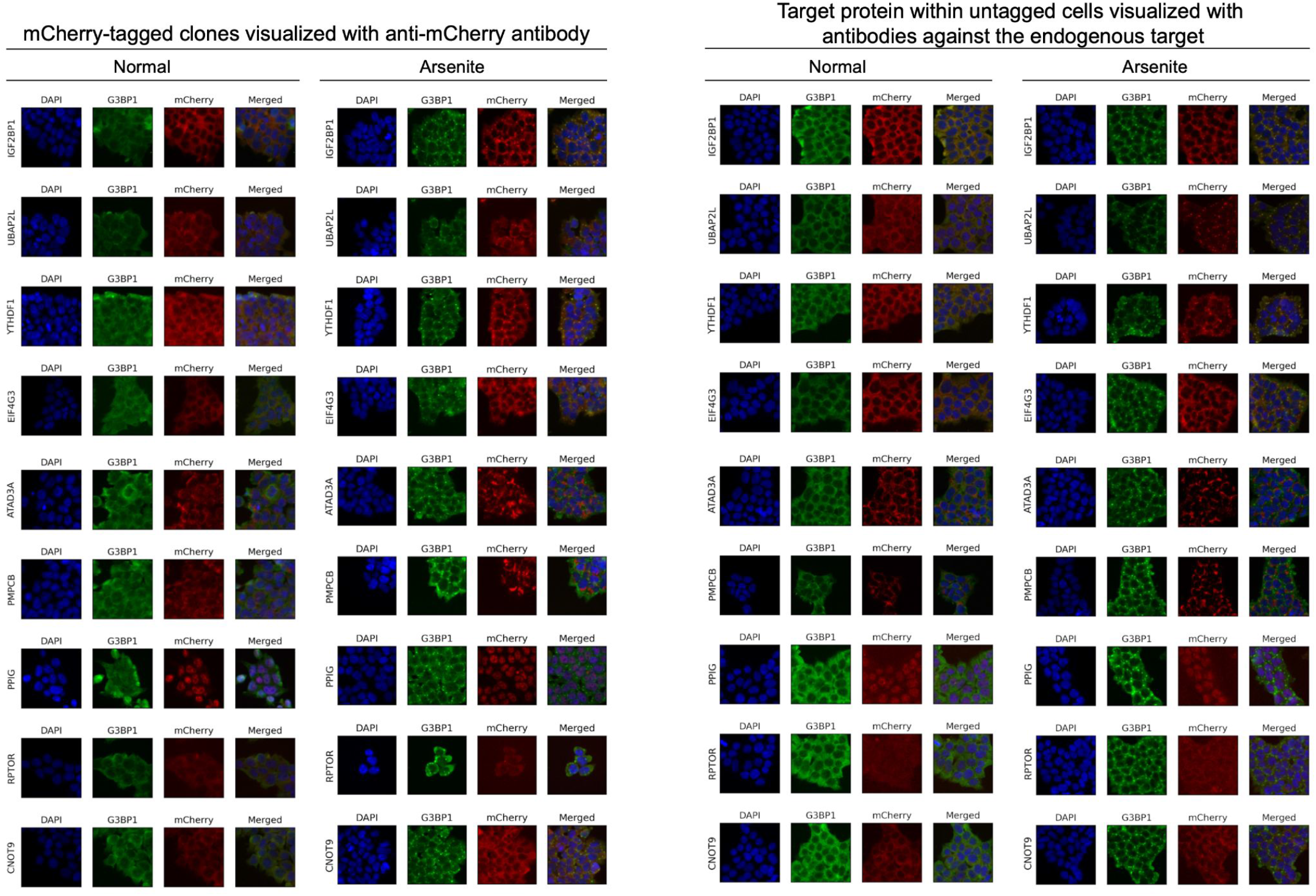
Comparison of protein localization and behavior between the mCherry-tagged version of a protein and the endogenous untagged counterpart. Images of stained cells under both homeostatic (normal) and stressed (arsenite) conditions are shown.

**Supplementary Figure 14.**
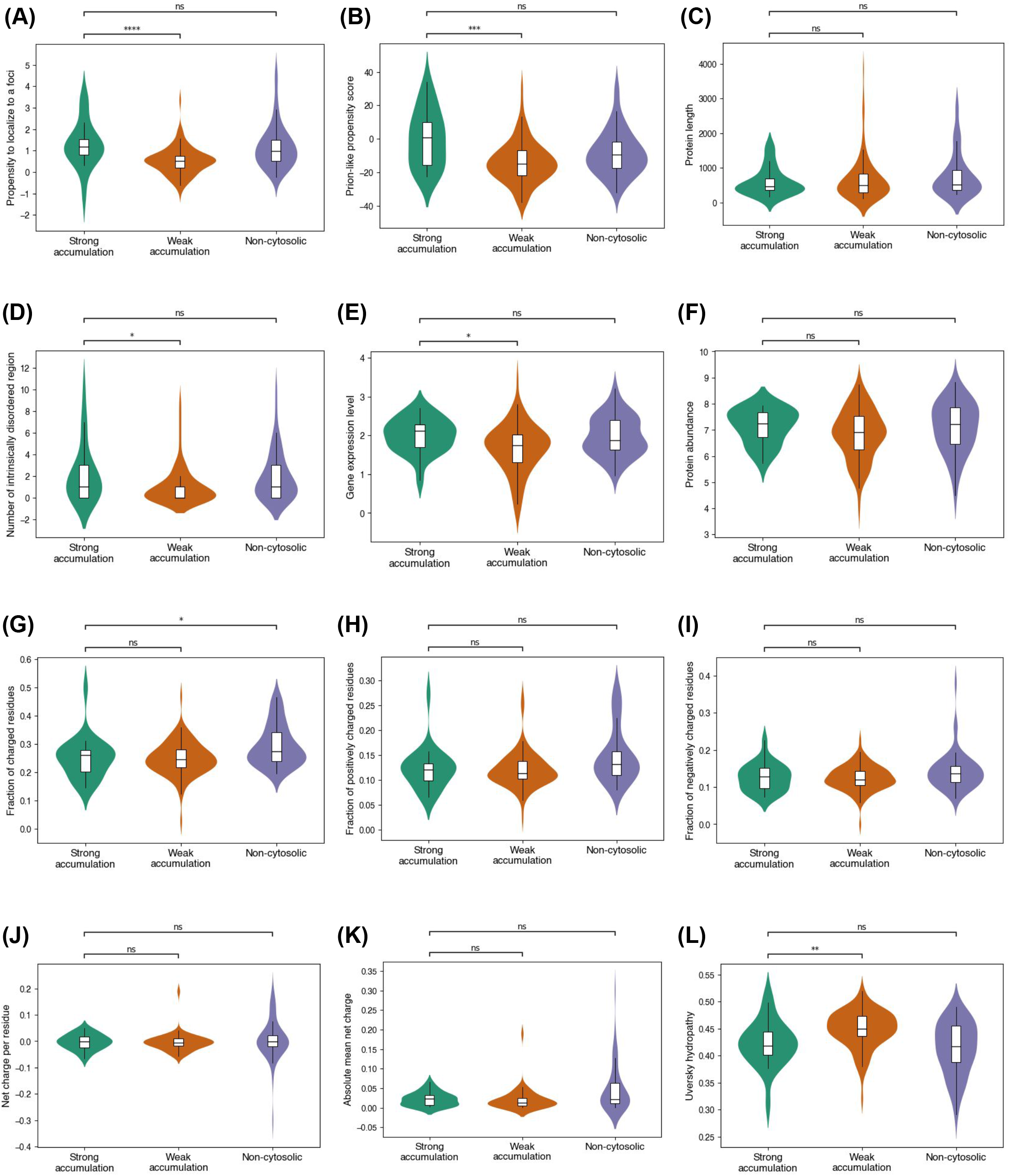
Analysis of the features which drive strong accumulation within SGs. Because all proteins that strongly gather in SG were primarily cytosolic analyses are done comparing cytosolic proteins that show strong vs weak accumulation. As an additional category, proteins that show non-cytosolic localization are also examined. (a) propensity to localize to a focus as determined by CatGranule, (b) prion-like propensity score as determined by Plaac, (c) Protein length (number of amino acids), (d) Number of intrinsically disorder regions, (e) RNA expression level represented as log (FPKM+1), (f) protein abundance extracted from mass spectrometry data, (g-i) fraction of charged residues, (j) net charge per residues at pH 7.4, (k) absolute mean net charge at pH 7.4, and (l) Uversky hydropathy were compared between the groups. (Mann-Whitney-Wilcoxon test, ns: not significant; * = p < 0.05; ** = p < 10^-2^; *** = p < 10^-3^; **** = p < 10^-4^)

## Notes

### Summary of Updates

Title and authors

## Reference

1. Bukhari, H. & Müller, T. Endogenous Fluorescence Tagging by CRISPR. Trends Cell Biol. 29, 912–928 (2019).

2. Palmer, E. & Freeman, T. Investigation Into the use of C-and N-terminal GFP Fusion Proteins for Subcellular Localization Studies Using Reverse Transfection Microarrays. Comp. Funct. Genomics 5, 742068 (1900).

3. Chudakov, D. M., Lukyanov, S. & Lukyanov, K. A. Fluorescent proteins as a toolkit for in vivo imaging. Trends Biotechnol. 23, 605–613 (2005).

4. Zhang, X. et al. Epitope tagging of endogenous proteins for genome-wide ChIP-chip studies. Nat. Methods 5, 163–165 (2008).

5. Nabet, B. et al. The dTAG system for immediate and target-specific protein degradation. Nat. Chem. Biol. 14, 431–441 (2018).

6. Stadler, C. et al. Immunofluorescence and fluorescent-protein tagging show high correlation for protein localization in mammalian cells. Nat. Methods 10, 315–323 (2013).

7. Cho, N. H. et al. OpenCell: Endogenous tagging for the cartography of human cellular organization. Science 375, eabi6983.

8. Reicher, A., Koren, A. & Kubicek, S. Pooled protein tagging, cellular imaging, and in situ sequencing for monitoring drug action in real time. Genome Res. 30, 1846–1855 (2020).

9. Schmid-Burgk, J. L., Höning, K., Ebert, T. S. & Hornung, V. CRISPaint allows modular base-specific gene tagging using a ligase-4-dependent mechanism. Nat. Commun. 7, 12338 (2016).

10. McCarty, N. S., Graham, A. E., Studená, L. & Ledesma-Amaro, R. Multiplexed CRISPR technologies for gene editing and transcriptional regulation. Nat. Commun. 11, 1281 (2020).

11. Xie, K., Minkenberg, B. & Yang, Y. Boosting CRISPR/Cas9 multiplex editing capability with the endogenous tRNA-processing system. Proc. Natl. Acad. Sci. 112, 3570–3575 (2015).

12. Hofmann, S., Kedersha, N., Anderson, P. & Ivanov, P. Molecular mechanisms of stress granule assembly and disassembly. Biochim. Biophys. Acta BBA – Mol. Cell Res. 1868, 118876 (2021).

13. Wolozin, B. & Ivanov, P. Stress granules and neurodegeneration. Nat. Rev. Neurosci. 20, 649– 666 (2019).

14. Doench, J. G. et al. Optimized sgRNA design to maximize activity and minimize off-target effects of CRISPR-Cas9. Nat. Biotechnol. 34, 184–191 (2016).

15. Song, B., Yang, S., Hwang, G.-H., Yu, J. & Bae, S. Analysis of NHEJ-Based DNA Repair after CRISPR-Mediated DNA Cleavage. Int. J. Mol. Sci. 22, (2021).

16. Thul, P. J. et al. A subcellular map of the human proteome. Science 356, eaal3321 (2017).

17. Van Nostrand, E. L. et al. A large-scale binding and functional map of human RNA-binding proteins. Nature 583, 711–719 (2020).

18. Kim, H. J. et al. Mutations in prion-like domains in hnRNPA2B1 and hnRNPA1 cause multisystem proteinopathy and ALS. Nature 495, 467–473 (2013).

19. Wall, M. L., Bera, A., Wong, F. K. & Lewis, S. M. Cellular stress orchestrates the localization of hnRNP H to stress granules. Exp. Cell Res. 394, 112111 (2020).

20. Suzuki, K. et al. In vivo genome editing via CRISPR/Cas9 mediated homology-independent targeted integration. Nature 540, 144–149 (2016).

21. Anzalone, A. V. et al. Programmable deletion, replacement, integration and inversion of large DNA sequences with twin prime editing. Nat. Biotechnol. 40, 731–740 (2022).

22. Pallarès-Masmitjà, M. et al. Find and cut-and-transfer (FiCAT) mammalian genome engineering. Nat. Commun. 12, 7071 (2021).

23. Ioannidi, E. et al. Drag-and-drop genome insertion without DNA cleavage with CRISPR-directed integrases. (2021) doi:10.1101/2021.11.01.466786.

24. Feldman, D. et al. Optical Pooled Screens in Human Cells. Cell 179, 787–799.e17 (2019).

25. Markmiller, S. et al. Context-Dependent and Disease-Specific Diversity in Protein Interactions within Stress Granules. Cell 172, 590–604.e13 (2018).

26. Youn, J.-Y. et al. High-Density Proximity Mapping Reveals the Subcellular Organization of mRNA-Associated Granules and Bodies. Mol. Cell 69, 517–532.e11 (2018).

27. Molina, R. S. et al. In vivo hypermutation and continuous evolution. Nat. Rev. Methods Primer 2, 36 (2022).

28. Coughlan, L., Cotter, P., Hill, C. & Alvarez-Ordóñez, A. Biotechnological applications of functional metagenomics in the food and pharmaceutical industries. Front. Microbiol. 6, (2015).

29. Walton, R. T., Christie, K. A., Whittaker, M. N. & Kleinstiver, B. P. Unconstrained genome targeting with near-PAMless engineered CRISPR-Cas9 variants. Science 368, 290–296 (2020).

30. Christie, K. A. & Kleinstiver, B. P. Making the cut with PAMless CRISPR-Cas enzymes. Trends Genet. 37, 1053–1055 (2021).

31. AU – Nageshwaran, S., et al. CRISPR Guide RNA Cloning for Mammalian Systems. J. Vis. Exp. e57998 (2018) doi:10.3791/57998.

32. Li, W. et al. MAGeCK enables robust identification of essential genes from genome-scale CRISPR/Cas9 knockout screens. Genome Biol. 15, 554 (2014).

33. Bolognesi, B. et al. A Concentration-Dependent Liquid Phase Separation Can Cause Toxicity upon Increased Protein Expression. Cell Rep. 16, 222–231 (2016).

34. Chu, X. et al. Prediction of liquid–liquid phase separating proteins using machine learning. BMC Bioinformatics 23, 72 (2022).

35. Lancaster, A. K., Nutter-Upham, A., Lindquist, S. & King, O. D. PLAAC: a web and command-line application to identify proteins with prion-like amino acid composition. Bioinformatics 30, 2501– 2502 (2014).

36. Holehouse, A. S., Das, R. K., Ahad, J. N., Richardson, M. O. G. & Pappu, R. V. CIDER: Resources to Analyze Sequence-Ensemble Relationships of Intrinsically Disordered Proteins. Biophys. J. 112, 16–21 (2017).

37. Szklarczyk, D. et al. The STRING database in 2021: customizable protein–protein networks, and functional characterization of user-uploaded gene/measurement sets. Nucleic Acids Res. 49, D605–D612 (2021).

